# Dynamics of complex genomic regions in giant genomes: MHC evolution in newts

**DOI:** 10.1101/2025.08.27.672527

**Authors:** W. Babik, K. Dudek, G. Palomar, M. Marszałek, G. Dubin, M. H. Yun, M. Migalska

## Abstract

Major Histocompatibility Complex (MHC) molecules are central to vertebrate adaptive immunity and MHC genes serve are key models in evolutionary genomics, offering insight into birth-and-death evolution, gene duplication, and the maintenance of genetic diversity. However, the organization and evolution of the MHC in species with giant genomes, such as salamanders, remain poorly understood. Here, we use comparative genomics, ontogenetic, tissue expression and polymorphism data across seven newt species to investigate MHC evolution in this group. Contrary to earlier suggestions of a massively expanded MHC in salamanders, we find that the core MHC region remains relatively compact, demonstrating that genome gigantism does not scale proportionally in this region. Our finding also challenges the model of coevolution between a single classical MHC-Ia gene and antigen processing genes (APGs), revealing instead several polymorphic and highly expressed putative MHC-Ia located at varying distances from the APGs. MHC-I genes exhibit lineage-specific duplications and signs of concerted evolution, resulting in poorly resolved phylogenies. In contrast, MHC-II genes are more conserved and exhibit extensive trans-species polymorphism. Expression and polymorphism patterns identify putative nonclassical MHC-Ib genes, likely repeatedly derived from MHC-Ia genes—paralleling patterns seen in mammals, but contrasting with the situation in fish and *Xenopus* frogs. In all seven species, some MHC-Ib genes show high relative expression during the larval stage but not at adulthood, suggesting a role in larval immunity. Our results underscore the importance of salamanders for understanding the evolution of complex regions in giant genomes and the architecture of the tetrapod MHC.

## Introduction

The Major Histocompatibility Complex (MHC) is a large genomic region originally identified as the genetic locus responsible for rapid allograft rejection in vertebrates — hence its name (Klein 1986). Subsequent research revealed that these incompatibilities are primarily caused by a subset of genes now known as the classical MHC genes, which encode molecules essential to the molecular-level self/non-self recognition and the initiation of the adaptive immune response (Murphy and Weaver 2016). Because of its critical role in transplant rejection and its associations with susceptibility to both infectious and autoimmune diseases, the MHC has been a subject of intense work. Decades of research have found it to be among the most polymorphic and structurally variable regions in the genome, shaped by a complex dynamics of host-pathogen co-evolution (reviewed in Radwan et al. 2020). Beyond immunological significance, the MHC has thus become a prime model in evolutionary and comparative genomics, offering insights into birth-and-death processes, concerted evolution, gene conversion, co-evolution of polymorphic genes, maintenance of genetic diversity, and immunogenetic trade-offs (Shiina et al. 2017; Kaufman 2018b; e.g., Hughes and Nei 1988, 1992; Migalska et al. 2019).

Classical MHC genes encode two types of cell surface glycoproteins (Fig. 1) that present peptide antigens to T cells: MHC class I (MHC-I, referred to as MHC-Ia) and MHC class II (MHC-II, Murphy and Weaver 2016; Pishesha et al. 2022). Class I molecules are ubiquitously expressed and present cytosol-derived antigens to cytotoxic CD8⁺ T cells. In contrast, class II molecules are primarily expressed on antigen-presenting cells and present extracellularly derived peptides to helper CD4⁺ T cells. In addition to classical MHC, there are also non-classical MHC genes (Adams and Luoma 2013). The non-classical MHC-I (MHC-Ib) molecules are typically non-polymorphic, have expression restricted to certain tissues, and often interact with innate immune cells or unconventional T cells (Mayassi et al. 2021). Some retain antigen-presenting capabilities, commonly for non-peptide antigens, and often assume roles at the interface of adaptive and innate immunity, or even outside the immune system entirely (for a comprehensive review in mammals, see Adams and Luoma 2013). The presence and function of MHC-Ib in non-mammalian vertebrates are poorly studied, but large, divergent families of MHC-Ib genes have been identified in fish (Grimholt et al. 2015), and several *Xenopus* frog MHC-Ib genes have been characterized in detail, demonstrating roles in tadpole immunity (Edholm et al. 2013, 2018). Non-classical MHC class II genes, such as DM, generally function as chaperones that facilitate peptide loading onto classical MHC class II molecules (Pishesha et al. 2022). DM genes are present in most tetrapods and lungfish but are absent in other fishes (Dijkstra et al. 2013; Almeida et al. 2020; Veríssimo et al. 2023).

**Fig. 1.**
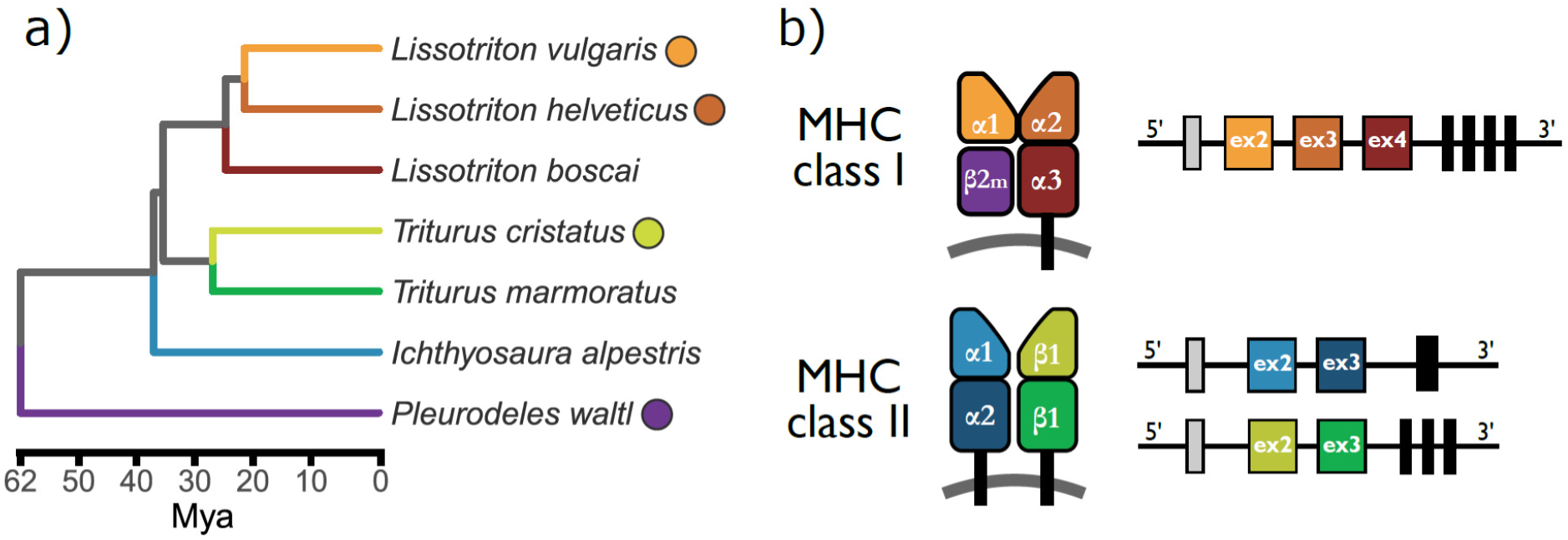
Newt phylogeny and a schematic representation of MHC genes and proteins. a) Time-calibrated phylogeny of the studied species (Stewart and Wiens, 2025); coloured dots next to the species name indicate the availability of reference genome. b) Schematic representation of the structure of MHC molecules and their genes. MHC class I molecules are composed of a heavy chain with three extracellular domains (α1, α2, and α3), paired with a non-variable β2-microglobulin (β2m) molecule, which is encoded by a gene outside the MHC region. MHC class II molecules are heterodimers, each consisting of an α and a β chain. Each chain contributes two extracellular domains: α1 and α2 for the α chain, and β1 and β2 for the β chain. The peptide-binding groove in MHC class I is formed by the α1 and α2 domains, whereas in MHC class II it is formed by the α1 and β1 domains. All MHC molecules also include a transmembrane domain and a cytosolic tail. There is a direct correspondence between structural domains and exons, as illustrated on the right-hand side. The first exon (grey box) encodes a leader peptide; each extracellular domain is encoded by a separate exon (indicated as “ex” with the exon number inside the box). The remaining regions—including the transmembrane domain and cytosolic tail—are encoded by a variable number of exons (black boxes), typically ranging from one to four, depending on the molecule type.

While nearly all jawed vertebrates studied to date possess MHC, its genomic architecture is highly variable. Efforts to reconstruct the primordial organization of the MHC and trace the evolutionary trajectories leading to the configurations observed in extant vertebrates remain an area of active research (Ohta et al. 2006; Flajnik and Kasahara 2010; Kaufman 2018b; Ohta et al. 2019; Veríssimo et al. 2023). One key insight from comparative studies is that the well-characterized genomic organization seen in mice and humans (the long-standing gold standard for MHC genomic research) — is actually a derived state, specific to placental mammals. In humans, the MHC spans 4-5 Mb, and contains well over 200 genes, many of which are not directly involved in immune response (Horton et al. 2004; Shiina et al. 2009). In humans — and more broadly in placental mammals — the Class I region, which includes MHC-Ia and some MHC-Ib genes, is separated from the Class II region by the Class III region. The Class III region contains both genes unrelated to immunity and immune-related genes with functions distinct from antigen presentation. The Class II region includes both classical and non-classical MHC-II genes, as well as antigen processing genes (APGs), which are critical for preparing and transporting peptides for loading onto MHC-Ia molecules. At the edges of the MHC locus lie the “extended MHC regions”, among which the extended Class II region appears more conserved across species. In contrast, the ancestral MHC organization is now widely accepted to consist of a core MHC-I/II region (hereafter referred to as “core MHC region”), where all essential components of the “adaptive MHC” (Veríssimo et al. 2023) are located in close proximity: MHC-I genes and their APGs, as well as MHC-II genes. This arrangement occurs across a wide range of non-eutherian vertebrates, including cartilaginous, sarcopterygian, and basal actinopterygian fish (Veríssimo et al. 2023), all three extant amphibian orders (He et al. 2023), lizards (Card et al. 2022), crocodiles (He et al. 2022), birds (Kaufman et al. 1999), monotremes (Zhou et al. 2021), and even marsupials (Belov et al. 2006).

The architecture of the MHC region may have far-reaching evolutionary consequences. Specifically, the physical proximity — or separation — of genes influences the frequency of recombination between them, thereby potentially promoting or constraining co-evolution among polymorphic genes that function together in a common pathway. This coevolutionary entanglement of interacting partners was proposed to limit their potential for gene duplication (Kaufman 1999; Ohta and Flajnik 2015; Martin and Kaufman 2022). A primary example is the co-evolutionary theory concerning APGs and MHC-I. In placental mammals, a proposed translocation event placed a large block of genes — the class III region — between MHC-I and its associated APGs (which remained with class II), disrupting their physical linkage. This structural change is thought to have lifted the co-evolutionary constraint, allowing for the emergence of a multigene MHC-Ia family and generalist APGs that provide multiple MHC-Ia molecules with broadly compatible, “average-best fit” peptides. However, the ancestral, tight linkage between MHC-Ia genes and their associated APGs, seen in non-eutherian tetrapods, has been proposed to promote co-evolution, while simultaneously constraining the expansion of MHC-Ia genes within the core MHC (for comprehensive reviews, see Kaufman 2015; Ohta and Flajnik 2015). Supporting this hypothesis, studies have found a single highly expressed MHC-Ia gene in the core MHC of both *Xenopus* and chicken, accompanied by more divergent clusters of MHC-Ib genes located further away — such as the *XNC* gene family in *Xenopus* and the *Rfp-Y* complex in chicken. This is at odds with growing evidence indicating that MHC-I in non-eutherian taxa often undergoes expansion through gene duplication — likely including MHC-Ia genes. Whether this expansion generally occurs within the core MHC-I region remains unclear (Minias et al. 2018, 2022).

Salamanders (Urodela), one of the three extant amphibian orders, constitute a group that could be particularly informative in shedding light on many aspects of the MHC evolution, including those described above. As an early branching tetrapod lineage, salamanders are a natural choice for exploring the evolution of MHC architecture across the vertebrate phylogeny. Notably, they are known to frequently possess duplicated MHC-I genes (Minias et al. 2022), making them good candidates for tests of APG-MHC-I co-evolution hypothesis. However, the exceptionally large genome sizes, ranging from 10 to 120 Gb (Gregory 2024), have long hampered genome-scale analyses in salamanders. Recent advances in sequencing technologies have yielded high-quality, chromosome-scale salamander genomes (Nowoshilow et al. 2018; Smith et al. 2019; Schloissnig et al. 2021; Brown et al. 2025). Although the complexity of the MHC region — and its rearrangements relative to the human — has introduced some confusion (for discussion, see Migalska et al. 2025), it is clear that we are now entering an era of unprecedented opportunity to resolve the MHC architecture in salamanders.

Apart from purely comparative aspects, such studies have a broad appeal, as several longstanding mysteries still surround salamander adaptive immunity. Similarly to other amphibians, they exhibit complex life cycles, often including an aquatic larval stage followed by metamorphosis (and typically a shift to a terrestrial lifestyle). Because of that, two major complications regarding the development of adaptive immunity emerge: 1) larvae face early pathogen exposure with limited time and cellular resources to develop full immune repertoires; 2) metamorphosis may necessitate a repeated self-tolerization of new, adult-type tissues. Research in *Xenopus* proposed a reliance on MHC-Ib molecules during larval stages as a solution (Edholm et al. 2013, 2018). However, since the ontogenic expression profiles of distinct MHC genes in Urodela remain largely unknown, it is unclear whether this mechanism could represent a universal feature tied to amphibian life history. A different molecular toolkit altogether could govern salamander immunity, as they have long been considered to exhibit subdued adaptive immune responses. Classical experiments reported slow (chronic) allograft rejection and weak mixed lymphocyte reactions in axolotls and several newt species, in contrast to anurans and other tetrapods (Cohen 1980; Pasquier et al. 1989; reviewed in Kaufman and Volk 1994). Over the years, various theories have emerged to explain these subdued adaptive immune responses, often suggesting deficiencies in MHC function — at one point even referring to it as an “Unrecognized MHC” (Kaufman et al. 1995). A detailed characterization of the structure and evolution of the salamander MHC would be a crucial first step toward resolving these controversies.

This work aims to present a comprehensive genomic and evolutionary view of the Major Histocompatibility Complex in newts (Salamandridae, Pleurodelinae). We investigate the genomic organisation of the MHC region, ontogenetic and tissue expression profiles, and the evolutionary history of the MHC gene family. To achieve this, we integrated publicly available chromosome-scale genome assemblies from four newt species with newly generated long- and short-read transcriptome data from multiple developmental stages and tissues across seven species (Fig. 1). Using comparative genomics, expression profiles, phylogenetic analysis, and structural modelling we trace MHC gene duplications, signatures of adaptive evolution, and identify putative classical and nonclassical MHC genes.

## Results

### Comparative genomics of the MHC region(s) in newts

We analyzed four publicly available newt genomes (*Lissotriton helveticus*, *L. vulgaris*, *Pleurodeles waltl*, *Triturus cristatus*), all assembled at the chromosome level. In each genome, there is a continuous MHC region assigned to a specific chromosome. Because the initial mapping of MHC transcripts to genomes indicated that automatic annotation of MHC genes was in many cases not accurate, MHC-I and MHC-II genes were annotated manually. The following description, based on this manual annotation, incorporates key observations informed by expression data and phylogenetic analyses, which are presented in detail later.

There are both considerable similarities and striking differences in the organization of the MHC region between the newt genomes (Figs. 2 & 3). A core MHC region, hereafter defined as spanning from the first MHC-II to the last MHC-I in the vicinity of APG or the last APG gene, is on an orthologous chromosome in all four species, though, as chromosome nomenclature is not standardized across assemblies, it is designated chr5 in *L. helveticus* and *L. vulgaris*, chr1 in *T. cristatus*, and chr6 in *P. waltl* (Fig. 2). In *Lissotriton* and *Pleurodeles* the MHC region is located at the terminal part of the chromosome reaching ca. 0.5 (*L. vulgaris*) – 9 (*P. waltl*) Mb from the end of chromosome (Fig. 2). In *T. cristatus* the MHC region is at the distance of > 120 Mb from the end of the 2.5 Gb chromosome. Notably, the orientation of the region differs between assemblies, with the extended class II region located terminally/distally in *L. vulgaris* and *T. cristatus* and internally/proximally in *L. helveticus* and *P. waltl*. The availability of linkage maps of *L. vulgaris* and *T. cristatus* (France et al. 2024) allowed us to establish that the MHC region is actually located at the opposite ends of linkage group (LG) 1 in these two species, and a comparison with *P. waltl* genome suggests that the region was translocated into a new location in *T. cristatus* lineage. The length of the core MHC region varies from 5.5 Mb in *P. waltl* to 16.7 Mb in *L. vulgaris*.

**Fig. 2.**
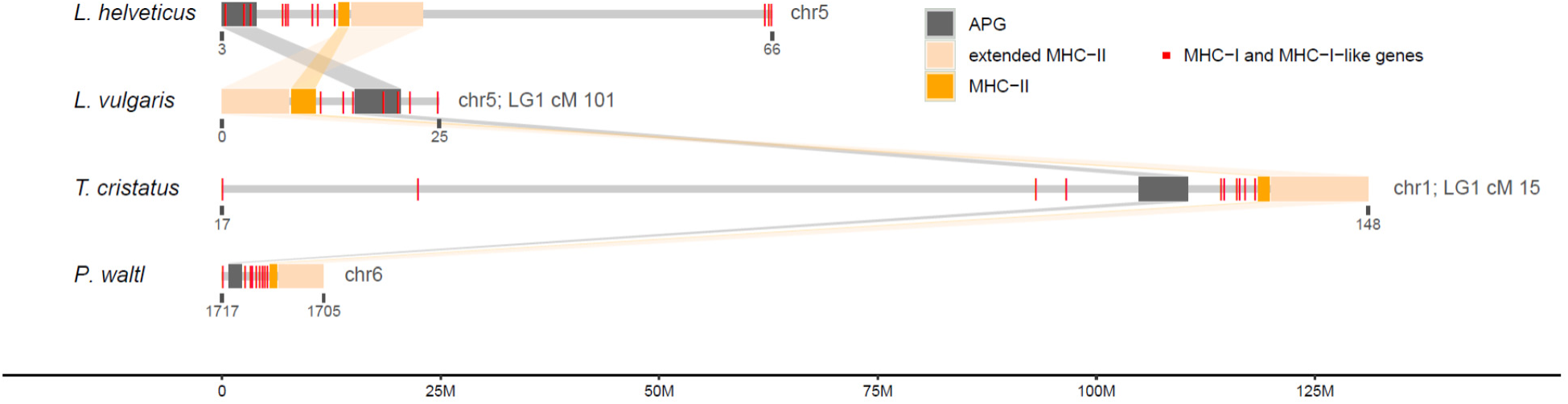
The MHC region in newt genome assemblies and on linkage maps. The location and length of the MHC region comprising APG, MHC-II, extended MHC-II gene and all manually annotated MHC-I genes located within 100 Mb. Note, that although in all genomes the MHC region is on the same chromosome, naming and coordinates follow the original genome assemblies. Comparative linkage maps of *L. vulgaris* and *T. cristatus* (France et al. 2024) indicate that the MHC region is located on the opposite ends of LG1 in these species – cM 101 and 15, respectively.

**Fig. 3.**
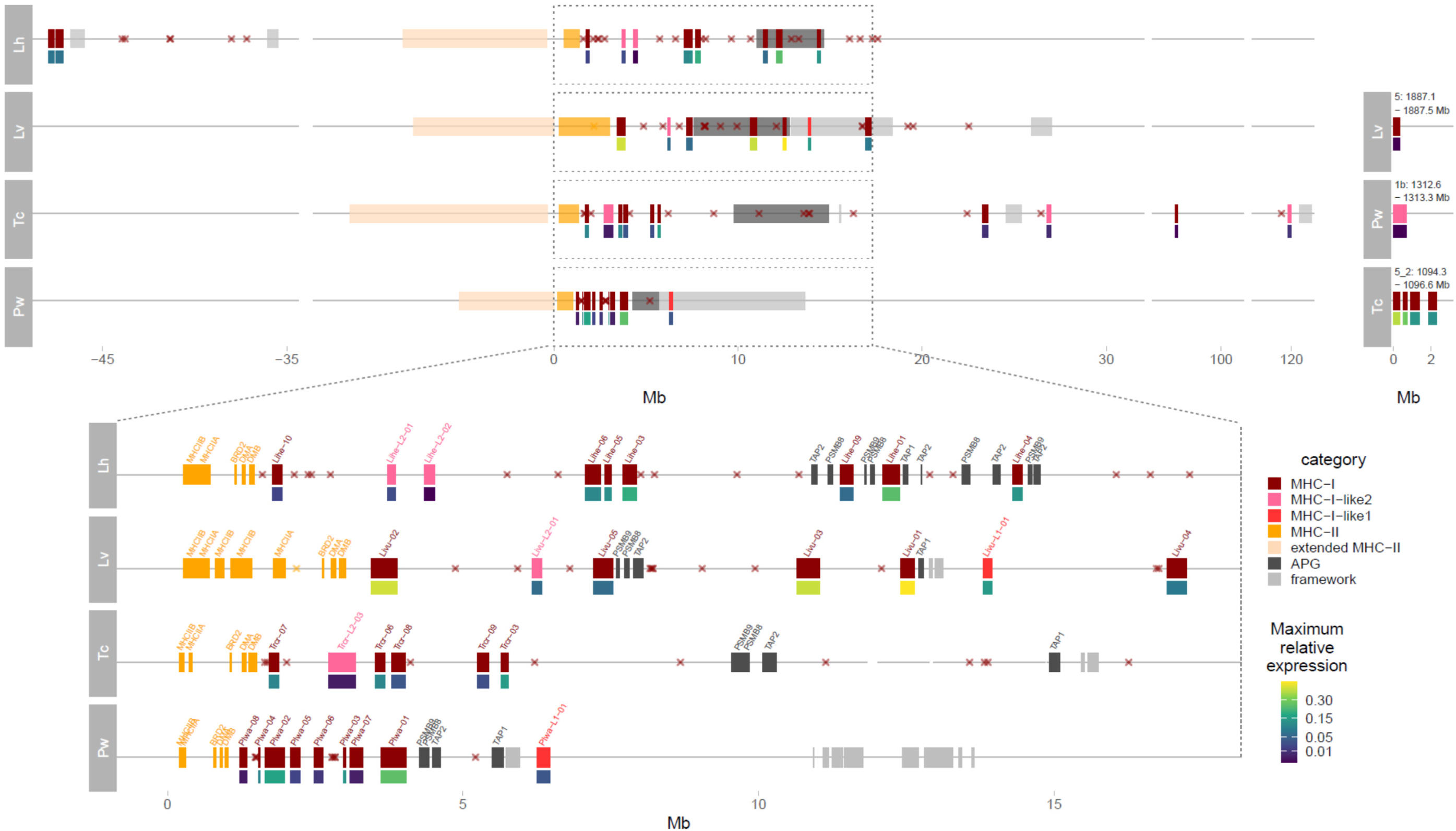
Genes in the MHC region(s). Upper part left – genes in the genomic regions corresponding to those in Fig. 2; APG, MHC-II extended class II and framework genes are indicated as colored blocks; MHC-I genes are shown individually, the width of the rectangles indicates gene lengths; filled rectangles below MHC-I genes show the maximum relative expression. Coordinates were rearranged so that the start/end of *COL11A2*, the extended class II gene adjacent in genomes to class II is given coordinate 0. Pseudogenes are marked with asterisks. Upper part right – MHC-I/MHC-like (if any) genes located outside the main MHC region. Lower part – the zoomed in “core” MHC region indicated with dashed rectangles in the upper plot. MHC-I, MHC-II and APG are marked and labelled individually with the width of rectangles corresponding to gene lengths. Maximum relative expression – the maximum fraction of total MHC-I expression (FPKM) attributable to a sequence across all RNAseq libraries from postmetamorphic individuals within a species.

In all species multiple MHC-I genes (from 3 in *L. vulgaris* to 8 in *P. waltl*), at least some of them highly expressed, are located between the MHC-II region and APGs (Fig. 3). In *L. helveticus*, most APGs are duplicated with four *TAP2*, three *PSMB8*, and two *PSMB9* copies, while in three other species each APG is a single copy gene. In both *Lissotriton* genomes some highly expressed MHC-I genes are interspersed among APG. Newt genomes contain also two divergent lineages of MHC-I sequences, which we name MHC-I-like1 and MHC-like2 (Fig. 4). In both genomes where we found it (*P. waltl* and *L. vulgaris*), MHC-I-like1 gene is close to *TAP1*, and MHC-I-like1 pseudogene in *T. cristatus* genome is also in this location. A single or two MHC-I-like2 genes are located between MHC-II and APG in *Lissotriton* and *Triturus* but on a separate chromosome in *Pleurodeles*.

**Fig. 4.**
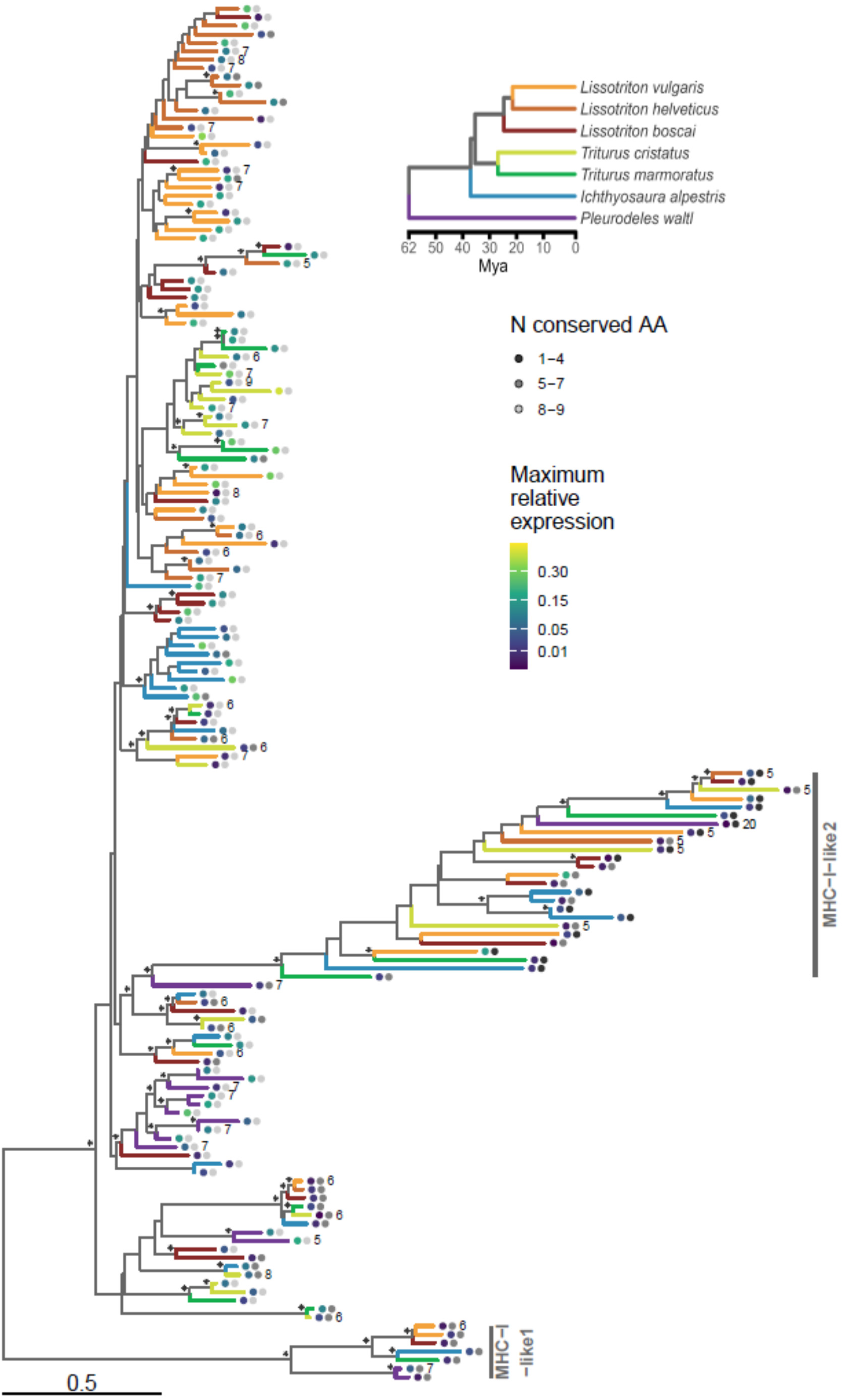
Phylogeny of MHC-I and MHC-I-like sequences. The RAxML-ng maximum likelihood tree constructed from protein sequences under the JTT+G4 amino-acid substitution model. The tree was rooted with *Andrias* sequence (GenBank AGY55973, not shown). Clades with the minimum bootstrap support of 70% (100 bootstrap replicates) are marked with asterisks. The tree contains sequences from the G+T dataset (as described in Materials and Methods) obtained from PacBio IsoSeq, de novo assembly of RNAseq and protein sequences predicted for manually annotated MHC genes in genome assemblies of *L. helveticus*, *L. vulgaris*, *P. waltl* and *T. cristatus*. Sequences predicted in genomes are marked with numbers indicating the number of protein-coding exons. Inset shows time-calibrated phylogeny of the studied species (Stewart and Wiens 2025). MHC sequences are color-coded according to the species. Maximum relative expression – the maximum fraction of total MHC-I expression (FPKM) attributable to a sequence across all RNAseq libraries from postmetamorphic individuals within a species. N conserved AA – the number of residues important for anchoring the termini of antigenic peptides containing amino acids that are conserved in classical MHC-I of most taxa (tippoint on the right).

In all species, some MHC-I genes are located outside of the core MHC region. Notably, MHC-I genes may be located on the same chromosome but > 40 Mb (*L. helveticus*) or even > 100 Mb (*T. cristatus*) from the core MHC region (Figs. 2 & 3). In three species, there are some MHC-I genes at the opposite end of the MHC chromosome (*L. vulgaris*), or on other chromosomes (*P. waltl* and *T. cristatus*, Fig. 3). Markedly, in *T. cristatus*, there are four highly expressed MHC-I genes that occupy ca. 3 Mb on chr5, not accompanied by any other genes usually found in the MHC region (Fig. 3). In the absence of another *Triturus* assembly or cytogenetic FISH evidence we cannot completely rule out the possibility that the apparent translocation is an assembly error. However, genes flanking the additional MHC region from both sides map to a continuous genomic region in other newt assemblies, rendering mis-assembly unlikely.

In all four genomes, MHC-II genes are in the same order (Fig. 3): *MHCIIB-IIA-BRD2-DMA-DMB* and span from <1 Mb (*P. waltl*) to > 3 Mb (*L. vulgaris*). The extended class II region is immediately adjacent to MHC-II in all species and spans from ca. 5 Mb in *Pleurodeles* to more than 10 Mb in *T. cristatus* (Fig. 3).

Finally, we examined immediate genomic context of the core MHC region, to see whether gene categories neighboring the MHC class I and II regions in mammals can be found in this group. Several framework genes (see Materials and Methods) were found in the vicinity of MHC in some species (Fig. 3). These are non-MHC genes interspersed in the eutherian MHC class I region (Amadou 1999), which are useful reference points for comparative analysis in mammals, but not necessarily beyond (Belov et al. 2006). Additionally, some MHC class III genes were found within or close to the core MHC genes in *T. cristatus*, though we note that precise localization of Class III is beyond the scope of the present article.

### Diversity of MHC sequences in newts

The MHC gene family frequently undergoes rapid expansions and contractions, leading to variation in gene content and blurred orthology among closely related species and even between haplotypes within species. As a result, a single reference genome per species is unlikely to capture the full extent of MHC diversity. Comprehensive sampling – both across individuals within species and across the phylogeny – is therefore key to understanding the evolution of MHC genes. To assess the diversity of MHC sequences in newts we complemented the sequences annotated in genomes (hereafter referred to as dataset G) with full-length coding sequences obtained from transcriptomes of altogether seven species (Table 1, S1, S2, Fig. 4). In addition to the four species with available genome assemblies, we included transcriptomic data from *Ichthyosaura alpestris*, *L. boscai* and *T. marmoratus*. The coding sequences were obtained mostly through PacBio IsoSeq, supplemented by a selection of well-supported transcripts assembled from Illumina RNAseq data. Translated sequences were clustered within each species to reduce redundancy (details are in Materials and Methods) and the longest protein from each cluster was selected as cluster representative. This set of combined genome- and transcriptome-derived sequences will hereafter be referred to as dataset G+T.

**Table 1.** The number of individuals at various developmental stages for which RNAseq data were obtained. Note that, for many individuals, libraries from more than one body part/tissue were sequenced (see text and Table S2). Developmental stages according to Shi et al. (1995): Young larvae - 38-44, Older larvae - 45-47, Metamorphosis - 48-49, Metamorph - 50. If offspring of more than one female was analyzed, numbers are separated by the ‘+’ sign.

| Species | Young larvae | Older larvae | Metamorphosis | Metamorph | Adult |
| --- | --- | --- | --- | --- | --- |
| *I. alpestris* | 2 + 2 | 1 + 1 | 1 + 2 | 1 + 1 | 2 |
| *L. boscai* | - | 2 | 1 | - | 2 |
| *L. helveticus* | 1 + 1 | 3 + 3 | 2 + 2 | 2 + 2 | 2 |
| *L. vulgaris* | 2 + 2 | 1 + 1 | 1 + 1 | 1 + 1 | 2 |
| *P. waltl* | 2 + 1 | 1 + 2 | 2 | 1 | 1 |
| *T. cristatus* | 2 + 2 | 1 + 1 | 2 | 1 | 2 |
| *T. marmoratus* | 1 + 1 | 3 + 2 | 2 | 2 | 2 |

Phylogenies MHC-I and MHC-II from the dataset G+T are shown in Figs. 4, 5. The most striking feature of the MHC-I tree in Fig. 4 is the presence of two well-supported clusters of sequences separated from the rest of the tree by long branches. These clusters, MHC-I-like1 and MHC-I-like2 are characterized by large deletions in α1 and α2 domains, respectively (Fig. 6). MHC-I-like proteins, in particular MHC-I-like2, commonly lack conserved amino acids at the anchor residues – those important for anchoring the termini of antigenic peptides, that are conserved in classical MHC-I of most taxa (Fig. 4, 6). MHC-I-like1 sequences were detected in five species (*I. apestris*, *L. boscai*, *L. vulgaris*, *P. waltl*, *T. marmoratus*), in each most likely a single gene (Table 2). We did not detect MHC-I-like1 sequences in either transcriptome data or in genome assembly of *L. helveticus*, while in *T. cristatus* genome there is an apparent MHC-I-like1 pseudogene, which suggests the loss of functional MHC-I-like1 in these two species. MHC-I-like2 sequences, which, as long branches in the phylogeny indicate, evolve quickly, were detected in all species, and they do not cluster according to the species phylogeny (Fig. 4 & Table 2) The number of variants per individual indicates multiple genes in some species. The remaining sequences on the MHC-I tree did not generally form well-supported clusters though there was a tendency to group by species or genus (Fig. 4). Given that MHC domains have different functions, they may be subject to different evolutionary pressures, including positive/purifying selection and gene conversion. Therefore, we additionally reconstructed separate phylogenies for MHC-I α1-3 domains (Fig. S1). These domains constitute the extracellular portion of the molecule, with the α1–α2 forming the highly polymorphic peptide-binding groove, and α3 interacting with accessory molecules and stabilizing the structure (Fig. 1). Phylogenies of individual α1-3 domains did not depart substantially from the full length protein phylogenies, though branch lengths, also those leading to MHC-I-like1 and MHC-I-like2, were much shorter for α3 than for the other two domains. Additionally, in α3, the MHC-I-like2 clade was not strongly supported (Fig. S1).

**Fig. 5.**
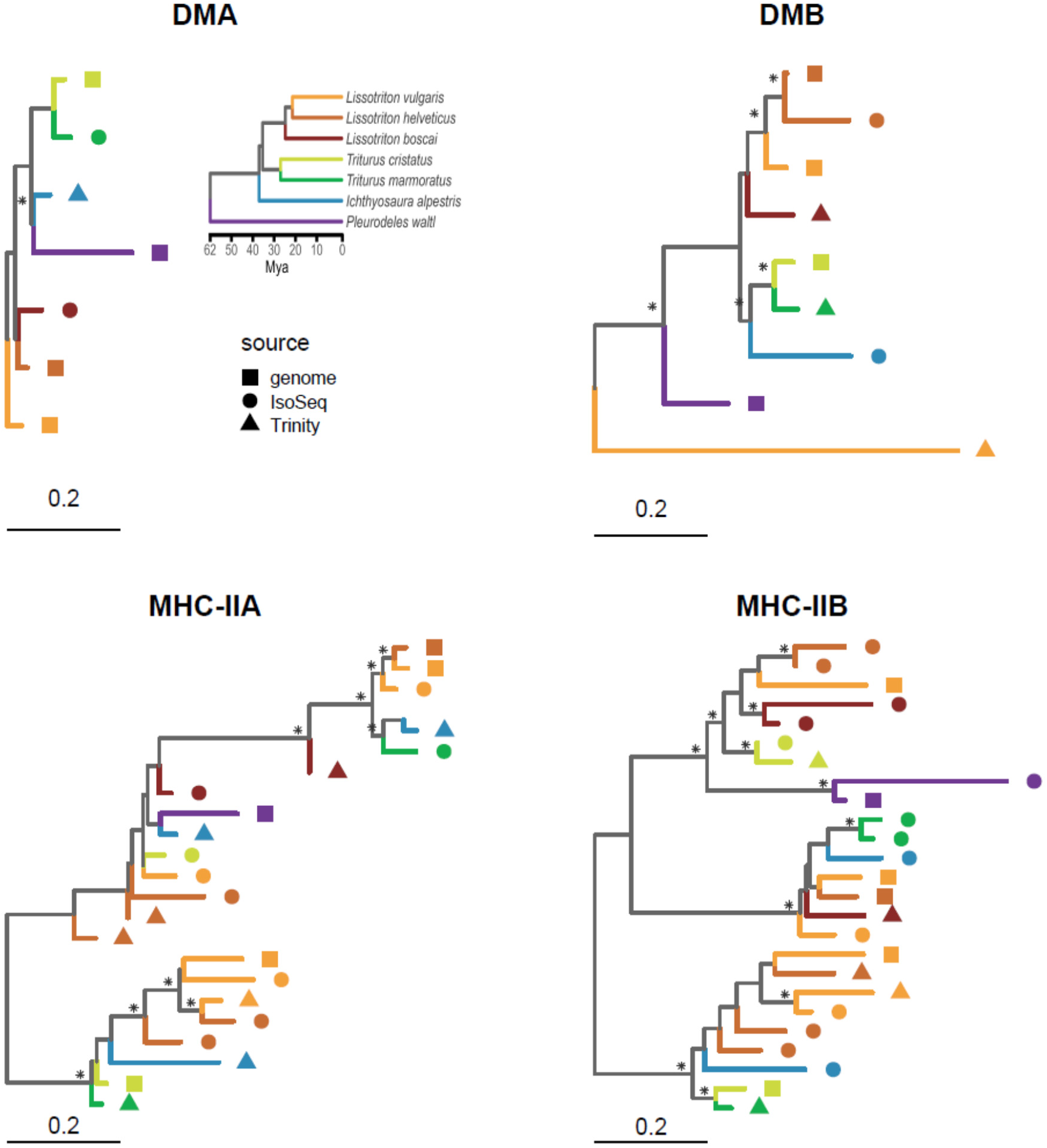
Phylogenies of MHC-II proteins. Separate trees were constructed for DM and other class II proteins as well as for α and β chains. The RAxML-ng maximum likelihood trees constructed from protein sequences under the JTT+G4 amino-acid substitution model. The trees were rooted with *Andrias* or axolotl sequences (not shown). Clades with the minimum bootstrap support of 70% (100 bootstrap replicates) are marked with asterisks. The tree contains sequences from the G+T dataset (as described in Materials and Methods) obtained from PacBio IsoSeq, de novo assembly of RNAseq and protein sequences predicted for manually annotated MHC genes in genome assemblies of *L. helveticus*, *L. vulgaris*, *P. waltl* and *T. cristatus*. Inset shows time-calibrated phylogeny of the studied species (Stewart and Wiens 2025). MHC sequences are color-coded according to the species.

**Fig. 6.**
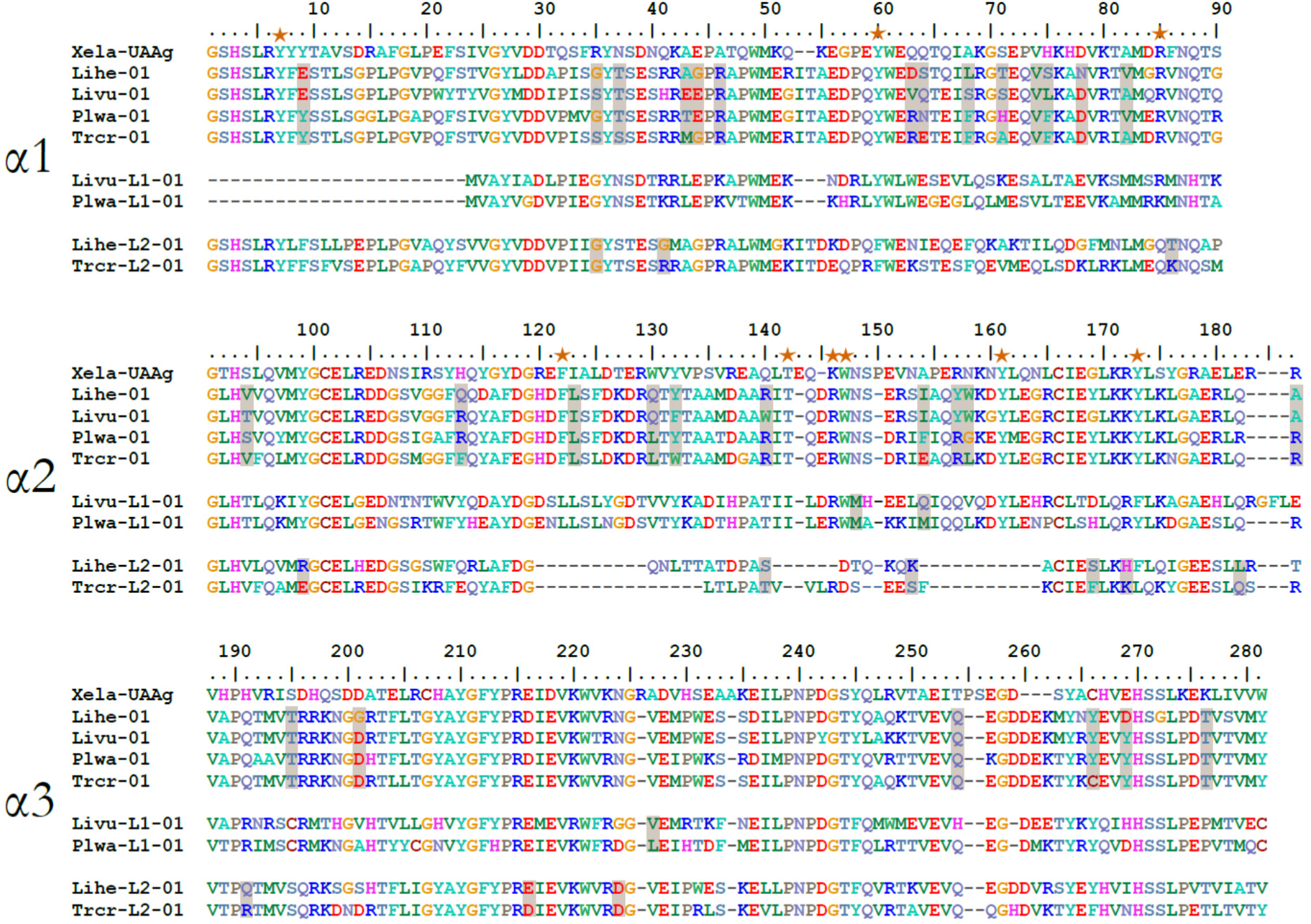
Protein alignments of α1-α3 domains, conserved and positively selected sites. Representative sequences of MHC-I, MHC-I-like1 (L1), and MHC-I-like2 (L2) genes from newt genomes were aligned to the *Xenopus laevis* allele with resolved crystal structure (Xela-UAAg, PDB: 6A2B). Positively selected sites identified in newts by FUBAR are shaded in grey; conserved anchoring residues are marked with orange stars. Species abbreviations: Lihe – *L. helveticus*, Livu – *L. vulgaris*, Plwa – *P. waltl*, Trcr – *T. cristatus*.

**Table 2.** The number of MHC-I genes and pseudogenes. Sp – species, N - number of genes (gen) or transcript clusters (tran) from a particular category (MHC-I, MHC-I-like1, MHC-I-like2); N pseudogenes refers to all categories combined. Transcript clusters were counted in the individual for which both PacBio Isoseq and RNA seq data were available, with the exception of *P. waltl*, for which these were counted based solely on IsoSeq data. NA – genome assembly not available; Total – total number of pseudogenes and gene fragments in the genome, in MHC – the number of pseudogenes and gene fragments in the MHC region as defined in Fig. 3, 10 Mb of MHC – the number of pseudogenes and gene fragments in the MHC region as defined in Fig. 3 and including flanking 10 Mb.

| Sp | N MHC-I |  | N MHC-I-like1 |  | N MHC-I-like2 |  | N MHC-I pseudogenes |  |  |
| --- | --- | --- | --- | --- | --- | --- | --- | --- | --- |
|  | gen | tran | Gen | tran | gen | tran | total | in MHC | 10Mb of MHC |
| *I. alpestris* | NA | 15 | NA | 1 | NA | 3 | NA | NA | NA |
| *L. boscai* | NA | 15 | NA | 1 | NA | 1 | NA | NA | NA |
| *L. helveticus* | 10 | 15 | 0 | 0 | 2 | 1 | 23 | 19 | 23 |
| *L. vulgaris* | 6 | 17 | 1 | 1 | 1 | 4 | 16 | 12 | 15 |
| *P. waltl* | 8 | 10 | 1 | 2 | 1 | 0 | 8 | 8 | 8 |
| *T. cristatus* | 11 | 12 | 0 | 0 | 3 | 1 | 14 | 14 | 14 |
| *T. marmoratus* | NA | 12 | NA | 1 | NA | 2 | NA | NA | NA |

The number of MHC-II sequences in the G+T dataset was much smaller than that of MHC-I, and their phylogenies showed simpler patterns. In case of non-classical MHC-II molecules, DMA phylogeny was poorly resolved, while DMB tree generally reflected species phylogeny, with one or two sequences per species (Fig. 5). Shorter DMA branch lengths indicate slower evolution compared to DMB proteins. Phylogenies of classical MHC-II, MHCIIA and MHCIIB, showed considerable similarity to each other – there was somewhat distinct *Pleurodeles* lineage and three divergent clusters, each of them containing sequences from all the remaining three genera. This pattern caused MHCIIA and MHCIIB phylogenies to depart drastically from species phylogeny. We did not find any evidence for the recently described MHC W type genes, that have both MHC-I and MHC-II features (Okamura et al. 2021), in newt genomes or transcriptomes.

### The number of MHC genes and pseudogenes

All studied newts have multiple MHC-I genes (Table 2). It is, however, not straightforward to estimate their number. Excluding MHC-I-like, there are from 6 (*L. vulgaris*) to 11 (*T. cristatus*) genes in the genome assemblies. The number of sequences retrieved from transcriptomes of individuals with IsoSeq data available (these estimates include both IsoSeq and Illumina RNAseq G+T datset sequences in these individuals) varied from 10 in *P. waltl* to 15 in *I. alpestris* and in all cases was larger than the number of genes detected in the assembly (Table 2). This is not surprising as divergent alleles found in heterozygous individuals could have been assigned into separate clusters. Also, the number of MHC-I pseudogenes and gene fragments differs between species, ranging from 8 in *P. waltl* to 23 in *L. helveticus*, with an overwhelming majority of pseudogenes and gene fragments within or in the vicinity of the MHC region (Fig. 3, Table 2).

In each genome, there are single DMA and DMB genes and, except *L. vulgaris*, single MHC-IIA and MHC-IIB genes. In *L. vulgaris* genome, there are two MHC-IIA and three MHC-IIB genes (Fig. 3). We did not identify DM pseudogenes in any genome, and generally, there were none or few MHC-II pseudogenes, the only exception being 21 MHC-IIA pseudogenes in *T. cristatus* genome. In contrast to MHC-I, MHC-II pseudogenes were located mostly outside the MHC region (Table S3).

### Ontogenetic and tissue expression profiles, polymorphism and identification of putative nonclassical MHC-I genes

Ontogenetic profiles of MHC expression can shed light on the timing of adaptive immunity onset in newts. Moreover, when combined with tissue expression patterns and polymorphism data, they can help to distinguish classical MHC-Ia from nonclassical MHC-Ib genes. Overall, changes in expression of MHC-I and MHC-II, were synchronized throughout development (Fig. 7). In *P. waltl* there is a steady increase of MHC expression through the larval stages, while in the remaining six species MHC expression in larval period and immediately after metamorphosis is at least an order of magnitude lower than in adults (Fig. 7), with a sharp increase visible only in mature individuals. The ontogenetic expression profile of MHC-I-like genes is very different – they are expressed at much lower level, and their expression remains low throughout the larval period, metamorphosis and in adults (Fig. 7).

**Fig. 7.**
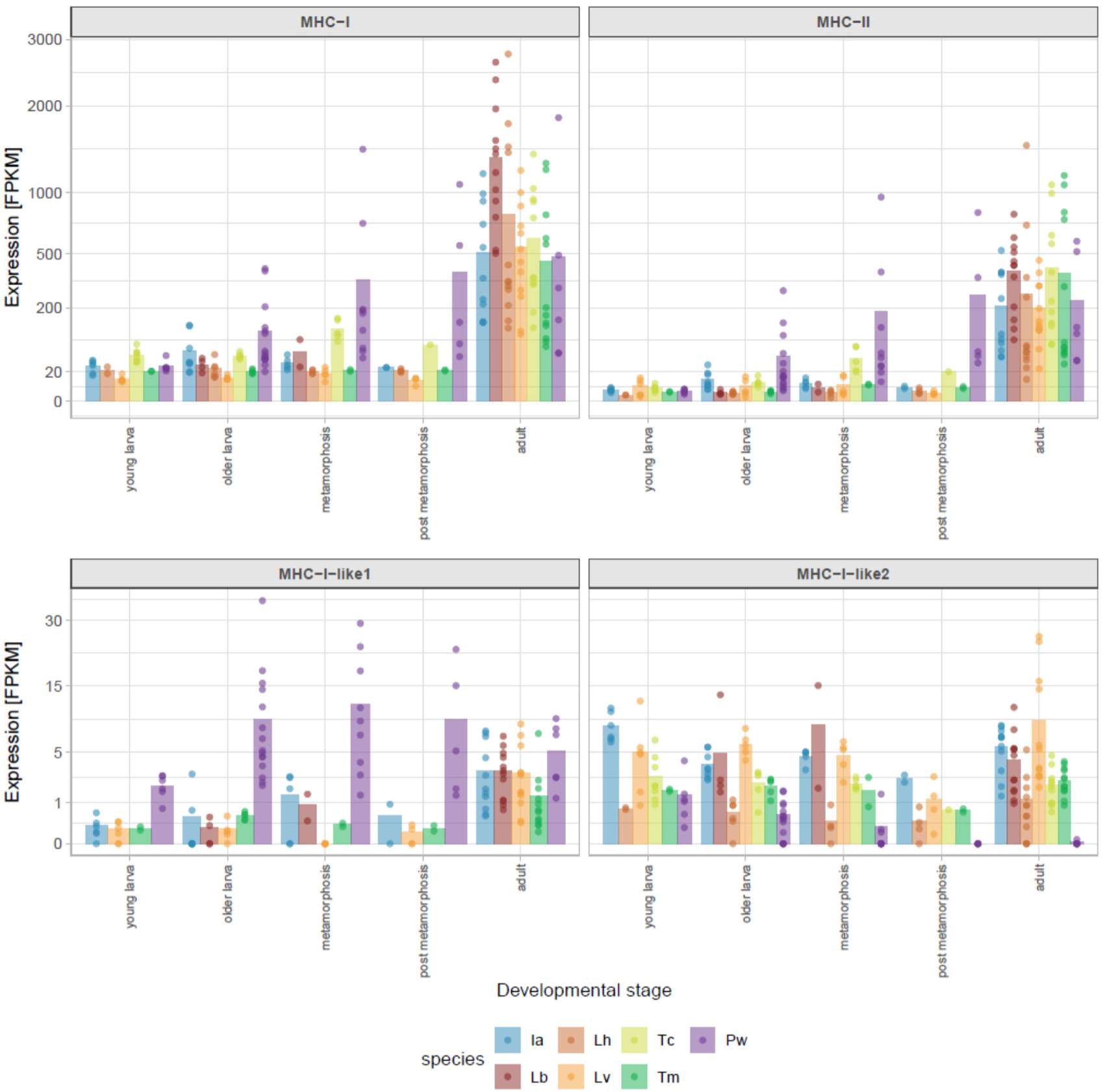
Overall MHC expression through ontogeny. Expression in Fragments Per Kilobase of transcript length per Million reads mapped (FPKM) of all MHC-I, MHC-II (MHC-IIA + MHCIIB), MHC-I-like1 and MHC-I-like2 sequences through the ontogeny. Points are expression values for RNAseq libraries (Table S2) and bars are averages for each species and stage. Species abbreviations: Ia – *I. alpestris*, Lb – *L. boscai*, Lh - *L. helveticus*, Lv - *L. vulgaris*, Pw – *P. waltl*, Tc – *T. cristatus*, Tm - *T. marmoratus*.

In both G and G+T datasets, only some sequences are highly expressed at any larval stage or in adults. While in *Triturus* and perhaps also *P. waltl* and *L. vulgaris* not more than two MHC-I genes are highly expressed in adults, in *I. alpestris*, *L. boscai* and *L. helveticus* several genes appear to have similarly high expression (Fig. 8 and S2). Moreover, many sequences highly expressed in at least one tissue/ontogenetic stage had conserved amino-acids in most (8 or 9 out of 9) anchor residues – one of the hallmarks of MHC-Ia molecules.

**Fig. 8.**
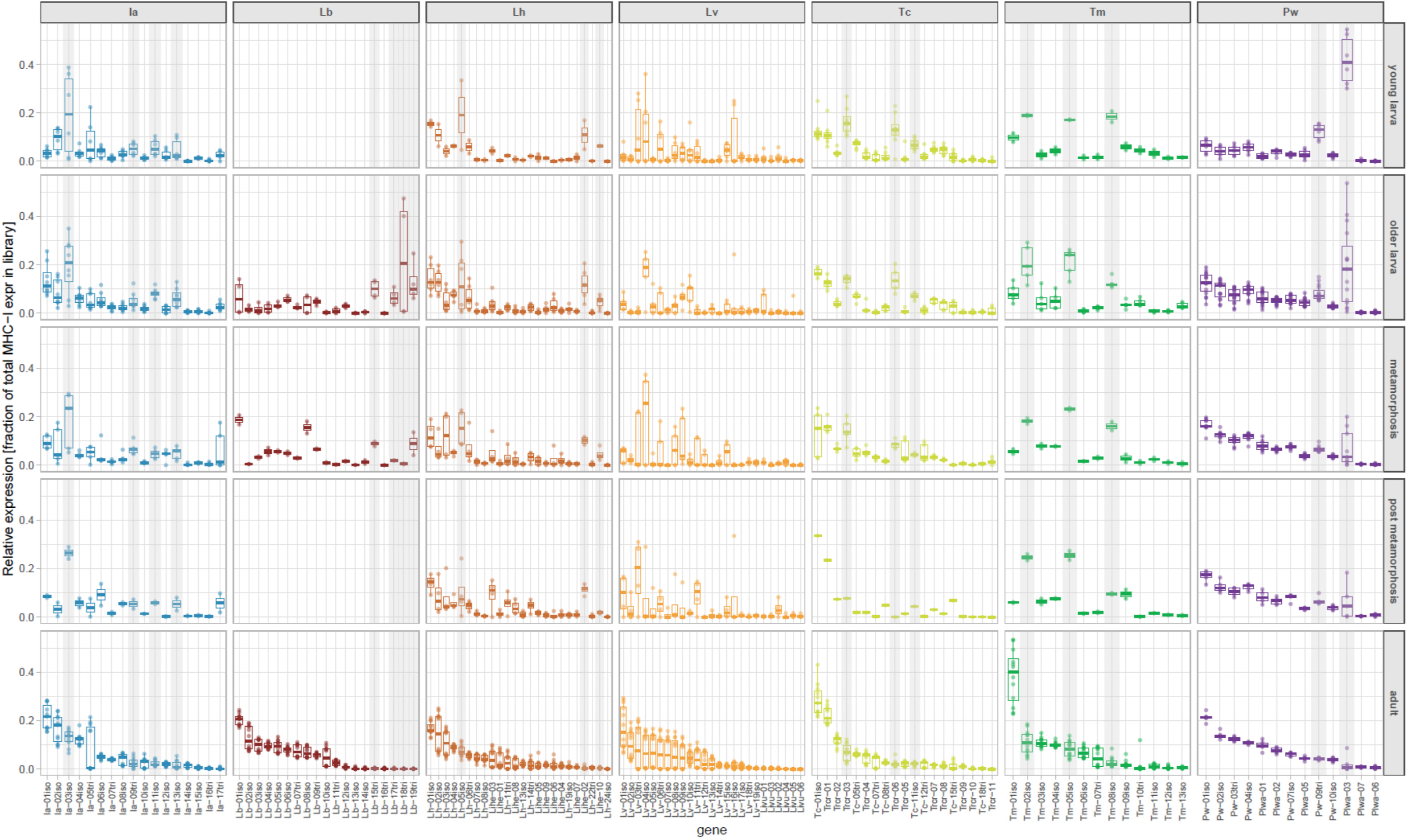
Expression of MHC-I sequences through ontogeny. Relative expression, i.e., FPKM for a particular cluster representative divided by the total MHC-I FPKM in a library. Boxplots show medians, interquartile and total ranges. Grey stripes indicate putative nonclassical (MHC-Ib) sequences. Species abbreviations: Ia – *I. alpestris*, Lb – *L. boscai*, Lh - *L. helveticus*, Lv - *L. vulgaris*, Pw – *P. waltl*, Tc – *T. cristatus*, Tm - *T. marmoratus*.

We did not detect pronounced differences between adult tissues in relative expression of particular MHC-I sequences, with the notable exception of *P. waltl Plwa-03* and *Plwa-04* which have much higher relative expression in skin (tailtip) than in other tissues (Fig. S3). The broad ranges of relative expression visible for some G and G+T dataset sequences in adults of all species except *P. waltl* (Figs. 8 and S2) reflect variation in mapping RNA-seq libraries coming from different individuals to these references. This behavior stems from underlying, inter-individual variation, suggesting that MHC haplotypes differ in gene content and carry divergent MHC-I sequences (Fig. S3).

Striking pattern of ontogenetic changes in the relative expression of some MHC-I genes was detected in all seven species. During the larval period, when the overall MHC expression is low, the relative expression of some MHC-I genes was much higher than in adults (Fig. 8 and S2). Such a pattern suggests a possible functional divergence from the role played by classical MHC genes and hints at a role in larval immunity or development. The sequences with high relative expression in larvae of *Triturus* and *Pleurodeles* are easily identified visually in Fig. 8 and S2, while the assignment of sequences to this category in other species is less straightforward. Still, such sequences do not form separate clades and occur in different parts of the phylogeny (Fig. S4). Therefore, regardless of difficulties in defining the exact set of such sequences, it is clear that they have emerged repeatedly and are probably undergoing fast turnover on evolutionary time scales.

For the four species with assembled genomes, we estimated polymorphism of MHC-I genes by mapping variants obtained in amplicon-based studies (Palomar et al. 2021; Gaczorek et al. 2023) to the genome and calculating the number of variants mapping to a particular MHC-I gene. Combining expression patterns with these polymorphism estimates (Fig. S5) suggests that sequences highly expressed in adults, which are also highly polymorphic, represent classical (MHC-Ia) genes. Sequences with high relative expression in larvae and lower expression in adults, which also show low polymorphism (Fig. S5), likely represent nonclassical (MHC-Ib) genes. The status of the remaining genes, which have low relative expression in all examined stages and tissues, as well as generally low polymorphism, is unclear. These may be additional MHC-Ib genes not involved in larval immunity or genes of less functional importance, perhaps on their way to pseudogenization.

Overall, in all species we see three broad categories of MHC-I genes: (i) putative classical MHC-Ia which expression increases in ontogeny, is very high in adults, and which exhibit high polymorphism; (ii) putative nonclassical (MHC-Ib) with low polymorphism, higher relative expression in larval stage and (in *P. waltl*) in tailtip tissue; (iii) either nonclassical or non-functional, with low expression throughout ontogeny and tissues, and more often than not low polymorphism. The distribution of these three categories on phylogeny indicates fast turnover and rapid evolution/diversification (Fig. S4).

### Evolution of MHC-I gene family in newts

The results presented above indicate a dynamic evolution of MHC-I family in newts with multiple gene duplications and losses. To obtain a more quantitative picture of this dynamics, we performed gene tree – species tree reconciliation in an attempt to infer the number of duplications and losses and identify clusters of orthologs. Reconciliation of the MHC-I gene tree with the species tree was performed separately for datasets G and G+T. The G dataset includes sequences of distinct genes, which should allow for a more reliable inference by avoiding confounding effects of allelic variation within a gene, but is limited to four species. The G+T dataset includes all seven species and many more sequences, but these originate from multiple individuals, and some may represent alleles of the same gene, potentially biasing the estimation of the number of gene duplications and losses. Each data set produced a single event history, for G 26 duplications, and 10 losses and for G+T 91 duplications and 66 losses. All species have experienced lineage-specific duplications and losses, though the exact numerical results in Table S4 should be treated with caution because some analyzed sequences may be derived from the same gene. However, we were not able to reliably reconstruct groups of orthologous sequences because, within the same event history, a large number of nodes in the reconciled tree could have been swapped without changing the overall number of duplications and losses.

Gene conversion may lead to the homogenisation of sequences of different members of gene families, resulting in concerted evolution that obliterates the signal of orthology. It is also an important mechanism generating new MHC variants by transferring small sequence segments between alleles and genes. To see whether concerted evolution via interlocus conversion could have contributed to poor resolution of MHC-I phylogeny and to the tendency for MHC sequences to group by species, we tested for gene conversion, estimated the length of conversion tracts, and checked whether some domains were involved in the process more often than others. For MHC-I-like1 and like2 which were analyzed together and with all species pooled, the signal of gene conversion was detected only for domain α3 and the tracts were shorter than in MHC-I.

In MHC-I, conversion was detected in all species, with a tendency to be more common in α3, which may contribute to the low sequence divergence seen in this domain (Fig. S1). We detected both conversion tracts contained within a single domain, i.e., within a single exon, and tracts spanning more than one exon. Because more of the latter were detected when mismatches within tracts were allowed (Table S5), and because tracts are expected to be relatively short so that they should not span across long newt introns, we suspect some signal may represent false positives. Nevertheless, the process of gene conversion appears widespread in newt MHC-I and may lead to concerted evolution of MHC genes within species.

### Selective pressures across MHC-I phylogeny

Phylogenetic analysis showed the presence of two divergent MHC-I-like lineages, in addition to MHC-I (Fig. 4). We were interested whether they experienced different selective pressures. In particular, we tested using RELAX and FUBAR: (i) whether a change in the strength of selection occurred along the long branches separating MHC-I-like1 and MHC-I-like2 from other sequences in MHC-I phylogeny, (ii) whether the number and location of codons under positive/purifying selection differ between the three categories of genes.

There was evidence of selection relaxation for MHC-I-like1: highly significant (p-values <0.001) RELAX test result for the branch leading to MHC-I-like1 cluster (k: 0.356, LRT: 401.7), and for the cluster itself (k: 0.476, LRT: 42.7), as compared to the MHC-I sequences. Accordingly, a direct, codon-based selection test (FUBAR) found only a few codons under positive or negative selection among MHC-I-like1 sequences (three and seven, respectively – posterior probability >0.9). The second divergent clade, MHC-I-like2, was associated with slight relaxation of selection intensity in branch leading to the clade (p-value: 0.021, k: 0.857, LRT: 5.3), but not in the cluster itself (p-value: 0.112). A direct selection test found 12 sites under positive and 50 sites under negative selection, which was considerably fewer compared to MHC-I sequences (30 and 122, respectively). Only two of the sites under positive selection overlapped between the two groups (Fig. 6). The selection tests were performed only on the three extracellular domains of MHC-I (α1, α2, α3), because the divergence of the remaining parts (i.e., the connecting, transmembrane and cytosolic portion of the molecules) rendered the alignment unreliable. Roughly half of the residues in the MHC-I were under negative selection, compared to ca. 20% in case of the MHC-I-like2 sequences. Most of the positively selected sites in MHC-I were located in the α1 and α2 domains (14 and 10, respectively), and majority exactly coincided or was immediately adjacent to residues forming peptide biding pockets as identified in *Xenopus* (Fig. 6). MHC-I-like2 sequences had much fewer positions under positive selection, scattered through the extracellular domains (four, six and three in α1, α2, and α3, respectively) (Fig. 6).

The overall concordance between the positively selected sites identified in the sequences forming the major MHC-I clade (Fig. 4), and the peptide binding pockets identified in *Xenopus* (Fig. 6) supports the classification of these sequences as the putative MHC-Ia (and perhaps some MHC-Ib). In contrast, markedly relaxed selection in MHC-I-like1 suggests a trajectory leading to pseudogenization, while in the case of MHC-I-like2, change in selective pressure strength and location of codons under positive selection hits to alterations in function, perhaps specialization.

### Structural modelling of MHC-I-like sequences

MHC-I-like1 and MHC-I-like2 sequences exhibit characteristic deletions within the α1 and α2 domains (Fig. 6), observed in transcripts and confirmed in the available genomes. MHC-I-like1 molecules lack a signal peptide and have a truncated α1 domain (missing 23 N-terminal amino acids). Lack of signal peptide likely precludes transport to endoplasmic reticulum, where MHC-I is folded and loaded with antigens (Paulsson and Wang 2003). Thus, this modification alone suggests departure from classical function. In contrast, MHC-I-like2 molecules possess a short N-terminal sequence resembling a signal peptide, an intact α1 domain, and substantial deletions within the α2 domain. To evaluate the structural impact of these alterations on the peptide-binding domains, we performed in silico modelling using AlphaFold 3. Presumably classical MHC-I proteins, included as references, were modelled with high confidence, exhibiting predicted local distance difference test (pLDDT) scores >90 in the α1–α2 domains. In comparison, the predicted α1–α2 domains of MHC-I-like molecules showed lower confidence, with overall pLDDT scores ranging from >70 to below 50 in the most uncertain regions. Most of the MHC-I-like1 models preserved the overall MHC-I fold, including the two antiparallel α-helices, but lacked the first β-strand and loop of the α1 domain (Fig. S6a), though one variant showed a closed conformation without open pockets (Fig. S6a, *L. vulgaris* conformation 2). Structural homology searches identified MHC and MHC-like molecules from other vertebrates as the closest matches, but none of the molecules contained deletions in the discussed region and thus the role of the deletion remains unknown. We speculate that a molecule with such an extensive deletion in the core would not fold altogether, but this speculation remains to be experimentally tested. In MHC-I-like2 molecules, deletions within α2 domain affect regions of the peptide-biding groove that, in classical MHC-Ia molecules, anchor the C-termini of bound peptides. Two β-strands forming the groove floor were either missing or significantly shortened, and a substantial portion of the α2 helix (including the conserved R146 and W147 residues) was also absent (Fig. S6b). Structural homology searches again primarily returned MHC and MHC-like molecules – these molecules however, lacked the discussed deletion. Interestingly, a notable exception was the viral MHC-I homolog m144 (from murine cytomegalovirus, PDB: 1U58), which shares similar deletions in the α2 domain. Unlike m144, which lacks a well-defined groove, the predicted MHC-I-like2 structures displayed seemingly an open conformation with a spacious, predominantly lipophilic pocket – this may however reflect the bias of AlphaFold towards classical MHC structures. Functionally, m144 is a molecular decoy mimicking host MHC-I, and expressed by the virus to prolong its survival by preventing natural killer cell activation (Natarajan et al. 2006). While this does not offer clues to the function of MHC-I-like2 in vertebrate hosts, it does support the plausibility of its folding pattern.

## Discussion

The results reported here advance our understanding of MHC evolution in salamanders and have broader relevance beyond this group. We will discuss first the broader implications of our work, i.e. the size and structure of the MHC region in giant genomes, coevolution of polymorphic genes within the MHC, and immunity of free-living early developmental stages of aquatic vertebrates. Then we will present insights more specific to salamanders, and conclude with a discussion of the limitations of the study and future prospects.

All studied newt species have a relatively compact core MHC region, which corresponds to the “adaptive MHC” of Veríssimo et al. (2023), although in some species (most notably in *T. cristatus*), additional MHC-I genes are present at a large distance or on other chromosomes. Nonetheless, the main class I (containing APGs) and class II regions are located next to each other and, unlike in eutherian mammals, are not separated by other genes. This presumably ancestral organization (Ohta and Flajnik 2015; Veríssimo et al. 2023) is thus retained both in salamanders and in frogs (Session et al. 2016; He et al. 2023). Importantly, our data do not confirm extensive expansion of the MHC region proposed in salamanders, or suggestions that the large-scale MHC architecture in this group may be more similar to that of eutherian mammals (Schloissnig et al. 2021). This earlier work was based on the analysis of synteny and emphasized the genomic location of genes and gene families found in MHC class III or linked to MHC class I in the eutherian mammals. As already demonstrated by Migalska et al. (2025), such an approach may indeed identify large genomic regions, which are, however, devoid of bona fide MHC genes. Thus, describing them as the MHC region is at best questionable. The situation is well illustrated by the *P. waltl* genome, where the genes found in the eutherian MHC region are located in two blocks at the opposite ends of chr 6, but the MHC genes are found only in a compact region within one of these blocks (Brown et al. 2025; the present study). The eutherian gene arrangement is derived, and the position of these genes should not be used to define the MHC region in other taxa. Our results thus demonstrate that the genomic gigantism in distantly related newts is accompanied by a relatively compact MHC region. Whether this holds in other taxa characterized by genomic gigantism is somewhat unclear. In another Urodele species with a huge (ca. 32 Gb) genome and resolved MHC architecture – the axolotl (*Ambystoma medium*) – the core MHC spans 13.8 Gb and has analogous organization to that found in newts (Migalska et al. 2025). In lungfish *Protopterus annectens*, the “adaptive MHC” region encompasses 20 Mb and is also relatively compact, considering the enormous genome size (>40 Gb). However, additional MHC-II genes are located at a considerable distance, therefore, if a more comprehensive definition were applied to define MHC region in the lungfish would span over 170 Mb (Veríssimo et al. 2023).

The compact genomic architecture is significant for a major theory positing that the tight linkage between classical MHC-I and APGs facilitates their coevolution, resulting in matching sets of MHC-I and APG alleles grouped in haplotypes (Joly et al. 1998; Kaufman 1999, 2015; Ohta and Flajnik 2015). Close proximity of coevolving polymorphic partners would minimize the formation of unfit MHC-I – APG recombinants. This theory, based on research in chicken and rat, assumes a single dominantly expressed MHC-I gene that coevolves with APGs (Kaufman 1999, 2015). Such a state has been suggested as ancestral for jawed vertebrates, with a “lucky” accident of a genomic rearrangement in mammals that broke the linkage between MHC-I and APG. This presumably led to the evolution of generalist APG, which in turn allowed duplication of classical MHC-I in mammals and an increased within-individual MHC variation (Kaufman 2018a). Notably, a key exception helped shape this hypothesis: in rats, a translocation of an MHC-Ia gene back into proximity with APGs (and loss of any remaining MHC-Ia genes from their Class I region) appeared to reinstate co-evolutionary dynamics (Joly et al. 1998). A major prediction of the coevolution hypothesis – a positive correlation between the MHC-I and APG diversity – was tested in a comparative framework using salamanders (Palomar et al. 2021). Salamanders commonly have more than one highly expressed MHC-I gene (Sammut et al. 1999; Fijarczyk et al. 2018; this study) and highly polymorphic TAP APG (Fijarczyk et al. 2018; Palomar et al. 2021). Palomar et al. (2021) found a positive correlation between the diversity of MHC-I and TAP genes (but not other APGs), indicating the coevolution between APGs and more than one classical MHC-I may be possible. The combined evidence from genomics, transcriptomics, and polymorphism presented here sheds further light on this matter. In three out of the four genome assemblies (*L. helveticus*, *L. vulgaris*, and *P. waltl*), the most highly expressed MHC-I gene is indeed the closest to the APG. However, the most highly expressed gene is not always the most polymorphic (although in all cases, polymorphism is substantial), nor is it necessarily the only highly expressed gene. *Lissotroton* newts simultaneously have multiple highly polymorphic genes, and two or more genes located in the proximity of APGs are highly expressed – both hallmarks of classical MHC-Ia genes. Taken together, these results indicate that more than one highly polymorphic and highly expressed MHC-I gene coexists with polymorphic TAP genes (Palomar et al. 2021), speaking against universal coevolutionary constraints forcing a single, dominantly expressed MHC-I gene.

Nonetheless, separation of APGs and MHC-I may indeed both facilitate MHC-I expansion and reduce TAP polymorphism – as exemplified by the exceptional situation found in *T. cristatus*. There, four highly polymorphic and highly expressed genes are located not in the core MHC region, where APGs reside, but on another chromosome. Remarkably, TAP diversity in seven *Triturus* species studied by Palomar et al. (2021), was among the lowest across salamander species and did not seem to correlate with MHC-I diversity (Fig. S7). This is consistent with an emergence of generalist non-polymorphic TAPs, which supply peptides to the classical proteins encoded by unlinked MHC-I genes. However, to further support this proposition, MHC organization in more *Triturus* genomes will need to be resolved. Overall, our findings point to a considerable plasticity in the genomic organization of the MHC-I in newts, which may translate into evolutionary flexibility in the interaction between key components of adaptive immunity.

Ontogenetic expression profiles of MHC genes, integrated with tissue-specific expression patterns and polymorphism data, allow (at least approximate) distinction between classical and non-classical MHC-I genes. All seven newt species examined have some MHC-I genes with high relative expression in larvae but not in adulthood. In the four species with available genomes, these genes also show low polymorphism. Both characteristics support their classification as nonclassical (MHC-Ib) genes, which could play a key role in larval immunity. Such a situation was described in *Xenopus* frogs, where XNC MHC-Ib restricts a population of unconventional T lymphocytes important in larval immunity (Edholm et al. 2013, 2018). However, unlike *Xenopus*, putative MHC-Ib genes in newts do not form a divergent gene family, but rather emerge repeatedly from classical MHC-I genes. This observation is in line with comparative data indicating that expanded families of putative, divergent MHC-Ib are not widespread among amphibians, and that the *Xenopus* genus might be a notable exception (He et al. 2023). These findings have broad implications for our understanding of developmental immunology in aquatic vertebrates. Such species must fight pathogen assault from early, free-living developmental stages, when constrained by the low number of available lymphocytes and the time needed to develop the diverse adaptive immune repertoire. The use of unconventional lymphocytes that carry T cell receptors of limited diversity could be an efficient solution in the face of such constraints (Edholm et al. 2014b; Robert and Edholm 2014). If such a strategy is indeed widespread, the rapid turnover of MHC-Ib genes, such as we see in newts, would suggest that populations of unconventional, innate-like T cells restricted by MHC-Ib molecules are ancient, and only MHC-Ib undergo rapid turnover, being repeatedly generated from classical MHC-I genes, in response to taxon-specific pathogen pressures. To test this hypothesis, ontogenetic profiling of lymphocyte populations across a wide range of aquatic vertebrates – using single-cell sequencing (Papalexi and Satija 2018; Rubin et al. 2021) – will be essential. We also note the evidence for the nonclassical status of genes reported here in newts comes from expression and polymorphism data, and does not have a strong, experimental support of *Xenopus* studies (Goyos et al. 2011; Edholm et al. 2014a). Therefore, complementary, functional studies at broader, phylogenetic scales will be essential, although – due to massive technical difficulties – likely not feasible in the near future.

In addition to the three themes of broad relevance discussed above, our results also shed light on several aspects of the developmental and evolutionary biology of MHC in salamanders. Our unique dataset tracks gene expression through larval ontogeny in newts, providing insights into the development of adaptive immunity in this group of amphibians. Particularly interesting is the contrast between *Pleurodeles* and the remaining six species, as for *P. waltl* substantial MHC expression appears already in larva and increases through metamorphosis. The onset of MHC expression (older larvae, stage 45-47 of Shi et al. 1995) coincides with the moment when thymectomy ceases to abrogate allograft rejection (Fache and Charlemagne 1975) – which marks the beginning of adaptive immune capacities. Experimental evidence for other newt species is modest, but some early work suggested the development of adaptive immunity may be generally more gradual in Urodeles, compared to Anurans (reviewed in Du Pasquier 1982). While this seems to be the case in *P. waltl,* poor overall larval MHC expression in other species contradicts this view. *Pleurodeles* is by far the largest of the taxa studied here, exceeding 75 g body mass, while *Triturus* weighs up to 15 g, *Ichthyosaura* 7 g and *Lissotriton* 5 g. It is therefore possible that species-specific body size (correlating with cell numbers, including lymphoid cells), rather than ontogenetic stage, determine the onset of adaptive immunity. This hypothesis can be tested with more data from salamanders that differ in body size.

Our data reveal several patterns in the evolutionary trajectories of immune genes in newts. The MHC-I gene family is remarkably dynamic, with abundant lineage-specific duplications, and some degree of concerted evolution through gene conversion resulting in poorly resolved phylogeny and a tendency of genes to cluster by species. Putative MHC-Ib genes originate repeatedly as discussed above, and numerous genes show both low expression and low polymorphism, indicating either nonclassical functions or the loss of functionality. Numerous MHC-I pseudogenes and gene fragments within the MHC region in all species testify to the dynamic birth and death evolution (Nei and Rooney 2005) at the temporal scale comparable to that in mammals or faster (Fortier and Pritchard 2024). The dynamic evolution of MHC-I contrasts with the situation in MHC-II family. Nonclassical DMA and DMB genes exhibit no signs of duplication and low polymorphism within species. There are also generally single classical IIA and IIB genes, with duplication detected only in *L. vulgaris*. The most remarkable feature of IIA and IIB genes is the presence of deeply divergent allelic lineages in all genera except *Pleurodeles*, suggesting long-term retention of transspecies polymorphism that may have persisted for over 30 million years. An ancient MHC category with both class I and class II features (MHC-W), which occurs in several fish lineages as well as in Hynobid and Ambystomatid salamanders (Okamura et al. 2021), was not found in any of the analysed species, suggesting their loss more than 60 million years ago. In addition, we identified two distinct lineages of MHC-I-like genes, separated by long branches from each other and from other MHC-I sequences. Expression of MHC-I-like genes starts in the early larva and stays similarly low throughout life. MHC-I-like1 is absent in *L. helveticus*, pseudogenized in *T. cristatus*, and shows signs of relaxed selection. It encodes a structurally atypical protein lacking a signal peptide and part of the α1 domain, making it unlikely to function in antigen presentation. In fact, it is unlikely that such protein folds at all in the α1 region. Structural modelling revealed a potentially artefactual cavity in the α1 domain, but we assume it was due to AlphaFold’s bias toward canonical MHC-I folds, on which the algorithm is trained. In contrast, MHC-I-like2 genes are present across all species, exhibit modest duplication, rapid evolution, and display a distinct distribution of positively selected sites in the α1 and α2 domains. Existence of a structurally analogous viral protein (m144) supports a functional potential, but the role of MHC-I-like2 is most certainly different from m144, and likely also diverges from the known MHC-I molecules. Definitive conclusions would require experimental structural data and functional assays.

The integration of chromosome-scale genome assemblies and transcriptomic data – particularly PacBio IsoSeq – has yielded valuable insights into the structure and evolution of the MHC in newts. However, both data types come with limitations. Assembling gigantic newt genomes remains a formidable challenge, making it unlikely that multiple high-quality assemblies will be available in the near future. While IsoSeq data can be obtained from multiple individuals and capture full-length transcripts, it does not provide information about the genomic location of those transcripts – a critical limitation given the complexity and rapid evolution of the newt MHC, including extensive copy number variation. Encouragingly, the relatively moderate length of the MHC region (in relation to the genome size) offers hope that advances in long-fragment DNA capture for long-read sequencing technologies (Iyer et al. 2024) will soon make it feasible to sequence full MHC haplotypes from multiple individuals. Such data are essential for a deeper understanding of the mechanisms driving MHC diversity. Further data on ontogenetic expression profiles from species that differ in body size are needed to test the hypothesis that body size determines the timing of the onset of adaptive immunity in salamanders. Finally, research on MHC genes should go hand in hand with characterising the populations of T cells and repertoires of T cell receptors through the ontogeny of diverse salamander species.

## Materials and methods

### Genome assemblies

We described MHC region(s) in chromosome-scale genome assemblies of *P. waltl* (Brown et al. 2025, Gen Bank Accession GCA_031143425.1) and three newt species sequenced and assembled by the Darwin Tree of Life Project: *L. helveticus* (GCA_964261635.1), *L. vulgaris* (GCA_964263255.1), and *T. cristatus* (GCA_964204655.1). The analyses were based on the primary haplotypes under the indicated accession numbers.

### Samples for transcriptome analysis

We analysed transcriptomes of the following newt species: *I. alpestris*, *L. boscai*, *L. helveticus*, *L. vulgaris*, *P. waltl*, *T. cristatus*, *T. marmoratus* (Table 1, S1, Fig. 1). Adult *P. waltl* from a laboratory colony in TU Dresden were mated, eggs and larvae developed at 19 °C and 12:12 L:D light regime. The following stages were sampled and preserved in RNAlater: young larvae (stages 38-44 Shi and Boucaut 1995), older larvae (stages 45-47), metamorphosing larvae (stages 48-49) and metamorphs (stage 50). Because *P. waltl* larvae are larger than those of other species, starting from stage 46, we sampled separately also intestine (middle part) and spleen. For all other species we sampled, during the breeding season, two presumably inseminated females from natural populations (Table S1). The females were kept in the laboratory until they deposited eggs, were then euthanized with MS222 and heart, intestine (middle part), liver, lungs, spleen and tail tissue were preserved in RNAlater. Eggs and larvae developed in the laboratory at room temperature under the natural light regime. Larvae were fed ad libitum with *Artemia* nauplii and then with live chironomid larvae. Larvae were euthanized in MS222 and preserved in RNAlater; depending on their size, they were cut into two (anterior/posterior) or three (head/middle/tail) parts, preserved and further processed separately (Table S2).

### RNA extraction and Illumina RNAseq

Total RNA was extracted from tissues preserved in RNAlater using RNAzol® RT (Sigma) or RNeasy kits (Qiagen). RNAseq libraries were constructed using Novogene Plant and Animal Eukaryotic Strand Specific mRNA (WOBI) approach and sequenced (2 x 150 bp) on the Illumina NovaSeq platform by Novogene.

### Full-length transcript sequencing

For *P. waltl* we used the available PacBio IsoSeq data obtained from the spleen RNA of the individual used for genome assembly (Brown et al. 2025). For the remaining six species, IsoSeq libraries from intestine (a tissue where diverse immune cells are abundant) RNA of a single individual per species were prepared and sequenced on the PacBio Revio platform by Macrogen. Note, that for each of these samples also short-read Illumina data were generated. PacBio HiFi reads were clustered into transcripts using SMRT Tools v. 13.1 lima, isoseq refine and isoseq cluster tools. Only high-quality (hq) IsoSeq transcripts generated by this pipeline were used in subsequent analyses.

### Identification of MHC transcripts

First, sequences of the axolotl MHC-I and MHC-II proteins were used as queries to identify MHC-I and MHC-II sequences in *P. waltl* reference transcriptome (Matsunami et al. 2019) using tblastn v. 2.15 (Camacho et al. 2009). Such identified MHC protein sequences were used to tblastn search the IsoSeq transcripts from all seven newt species at the E-value threshold of 1e-30. TransDecoder v 5.7.1 (Haas et al. 2013) was used to identify Open Reading Frames (ORF) in IsoSeq transcripts (both complete and partial ORFs were allowed, single best ORF per transcript). Because a large number of transcripts were identified (ca. 700 in the case of MHC-I) and visual inspection of their protein alignments indicated that sequences were often highly similar but differed by the presence/absence of entire exons, to reduce redundancy we clustered the proteins within each species based on sequence identity. Clustering was performed using Clusterize() from the R package DECIPHER (Wright 2016) at the protein sequence divergence threshold 0.1, ignoring regions containing gaps (penalizeGapLetterMatches = FALSE). The longest protein was selected as a representative of each cluster. Cluster representatives of the minimum length of 240 amino acids (aa) were aligned using DECIPHER::AlignSeqs(). The resulting alignments were visually inspected and sequences lacking more than half of any of α/β 1-3 domains, or showing extreme divergence in a part of the alignment and high similarity to at least one other sequence in the alignment were removed.

Because some IsoSeq clusters were represented by few HiFi reads and because some divergent MHC lineages contained representatives of only some species, we suspected that our IsoSeq dataset may have missed some MHC sequences, especially those poorly expressed in the intestine. Therefore we supplemented the IsoSeq dataset with sequences of transcripts obtained from de novo assemblies of short RNAseq reads. Separate assemblies were obtained for each of five individuals per species, including adults, metamorphs, and, for some species, also advanced larvae (Table 1, S2), reads from all individual’s RNAseq libraries were pooled and assembled with Trinity v. 2.15 (Grabherr et al. 2011). Trinity assemblies were searched for MHC sequences using tblastn with the axolotl and *P. waltl* MHC-I proteins and the axolotl MHC-II proteins as queries; sequences with hits of min. 90 aa alignment length and E-value < 1e-40 were retained.

TransDecoder was used to identify the single best ORF per Trinity transcript and the resulting proteins were clustered together with representative sequences of IsoSeq clusters. Although the majority of Trinity proteins fell into previously identified clusters, some new clusters which did not include IsoSeq sequences and had only Trinity sequences of at least 240 aa appeared.

Representative (longest) sequences from such clusters were visually examined for the signatures of chimaerism, and those that passed this check were added to the dataset. It is important to note here that Trinity sequences, resulting from de novo assembly of short reads, may not represent true MHC sequences because of the difficulties of reconstructing full-length transcripts of similar genes from multigene families from short reads. We decided to include such sequences because they represent, albeit imperfectly, extra MHC diversity present in individuals or tissues for which IsoSeq was not available or lowly expressed.

### The genomic organisation of the MHC region

The genomic location of the “core” MHC region and any disparate genome fragments that may contain MHC genes was established by blastn searches with cluster representatives obtained as described above. To annotate genes both within these regions and in their vicinity, we adopted two routes. Protein coding genes other than MHC-I and MHC-II were automatically annotated. We used the available gene predictions for *P. waltl* (Brown et al. 2025) and performed de novo prediction with BRAKER3 (Gabriel et al. 2024) for the remaining three genomes. Because de novo gene prediction is a complex and computationally intensive task, especially in large genomes, such as those of newts, we limited our prediction to the MHC region(s) (both core and disparate). We added other continuous genomic regions, totalling 950 Mb in *L. helveticus* and *L. vulgaris* and 700 Mb in *T. cristatus*, to reach a sufficient number of well-supported genes required to train prediction algorithms in BRAKER3. The available version of *T. cristatus* genome assembly is already softmasked for repeats, while in the case of *L. helveticus* and *L. vulgaris* assemblies, we performed repeat identification and softmasking using RepeatModeler v. 2.0.6 (Flynn et al. 2020) and RepeatMasker v. 4.1.7 (repeatmasker.org). Both RNAseq data from target species and vertebrate proteins from the OrthoDB database (Tegenfeldt et al. 2025) were used for gene prediction. Predicted protein sequences were extracted from the genome assemblies using gffread v 0.12.8. (Pertea and Pertea 2020) and annotated using eggNOG-mapper v2 (Cantalapiedra et al. 2021).

Because the initial mapping of cluster representatives to genomes indicated that automatic predictions of MHC genes were in many cases not accurate, MHC-I and MHC-II genes were annotated manually (also when present outside of the core MHC region). We used the following sources of information: i) gene models from automatic prediction, ii) mapping of all RNAseq libraries from a target species to the genome with hisat2 v. 2.2.1 (Kim et al. 2019), iii) mapping of all cluster representatives (obtained as described above) from the genus to the genome with minimap2 v 2.28 (Li 2018), iv) mapping of all IsoSeq transcripts from the species to the genome with minimap2. Tracks representing all the data were visualised in IGV v. 2.18.4 (Robinson et al. 2011), coordinates of coding sequences were identified and exported as .bed files. The retrieved sequences were checked for internal stop codons, and those that did not contain them were considered as putative MHC genes, forming the dataset G.

To identify putative MHC pseudogenes or gene fragments, we extracted for each species exons > 100 bp from all annotated MHC genes, and used blastn to search the genome at the E-value threshold of 1e-10 with scoring adjusted to divergent sequences (-reward 1 -penalty -1 -gapopen 3 -gapextend 2). Hits with at least 70% sequence identity covering at least 80% of the query sequence located outside manually annotated MHC genes were considered to represent pseudogenes or gene fragments. Complete annotations of the MHC genes and other genes in the MHC region are provided as .gff files in Dryad repository.

To ease comparative analysis, we classified all the annotated functional genes following categories used by Belov et al. (2006): (i) MHC Class I (putative classical or non-classical), (ii) MHC Class II (including classical MHC-II, DM, and a non-MHC, yet closely linked with Class II region *BRD2* gene), (iii) Antigen Processing Genes (APGs, which include *PSMB8*, *PSMB9, TAP1* and *TAP2*), (iv) Extended class II (from *COL11A2* to *ZBTB22* – if present, including *TAPBP*), (v) MHC Class III (if present), (vi) Framework region genes (conserved, non-MHC genes known to be interspersed in the eutherian MHC class I region (Amadou 1999), found in the vicinity of the core MHC region in non-placental mammals (Belov et al. 2006), and more dispersed in non-mammalian genomes). These categories are well suited to describe non-eutherian MHC organization while maintaining clear correspondence to the gold standard of the human MHC nomenclature. For clarity of visualisation, we show only synteny-informative Framework genes (i.e., present in more than one species) and omit genes that are not mapping to human MHC.

### MHC-I polymorphism

One of the key features distinguishing classical MHC genes from nonclassical is their high polymorphism. To assess polymorphism of the MHC-I annotated in the genomes we used amplicon sequencing data from previous studies (Palomar et al. 2021; Gaczorek et al. 2023). As a measure of polymorphism we used the number of MHC alleles mapping to a particular gene. This approach is approximate, as alleles may not always be most similar to the reference sequence of their genes of origin due to the high divergence between alleles and interlocus recombination, the genes of origin may be missing from the reference assembly, and alleles may map equally well to multiple genes. Still, even such an approximate measure should provide meaningful information about polymorphism. The available alleles were blastn-mapped to the respective genomes at the E-value threshold of 1e-30. Because the majority of alleles produced multiple hits, we considered as valid all hits within 0.9 of the bitscore of the best hit. Each hit was given the weight 1/n, where n was the number of hits, allowing an allele matching multiple locations in the genome to be counted as a fraction in each, avoiding overestimation of polymorphism for similar genes.

### Phylogenies of MHC proteins

Clustering of MHC proteins, following the procedure described above, was repeated to include also the proteins predicted in genomes. When available, the longest proteins predicted in the genome were used as cluster representatives. The sequences formed the G+T dataset. The G+T protein sequences, together with the axolotl or *Andrias* as outgroups, were aligned with DECIPHER::AlignSeq(), separately for MHC-I, MHC-IIA, MHC-IIB, MHC-DMA, MHC-DMB, as well as for MHC-I α1, α2, and α3 domains. The amino-acid evolution models were identified by ModelTest-NG v. 0.1.7 (Darriba et al. 2020); because for all alignments JTT + G4 or similar models were selected, we decided to use JTT + G4 model for all phylogenetic analyses. Maximum likelihood (ML) phylogenies were reconstructed in RAxML-NG 1.2.2 (Kozlov et al. 2019), and robustness of the obtained topologies was tested with 100 bootstrap replicates.

**MHC-I anchor residues** are defined as amino acids at nine residues important for anchoring the termini of antigenic peptides that are conserved in classical MHC-I of most taxa (Kaufman et al. 1994; Sammut et al. 1999; Almeida et al. 2021). The number of residues with conserved amino acids may be helpful in distinguishing classical and nonclassical MHC-I sequences and we calculated it for our sequences.

### MHC-I and MHC-II expression through ontogeny and across tissues

Expression was estimated by mapping RNAseq libraries to two types of references: (i) for the four species with the available genome assemblies, RNAseq reads were mapped to genomes with hisat2 and reads mapped to the annotated MHC genes, pseudogenes and gene fragments were counted with featureCounts v. 2.0.8 (Liao et al. 2014) – this analysis provided detailed expression information for MHC sequences present in reference genomes, (ii) for all seven species, RNAseq reads were mapped to the cluster representatives using Bowtie2 v. 2.5.1 (Langmead and Salzberg 2012) and the number of reads mapped to each reference was calculated with samtools idxstats – this analysis provided expression estimates for all full length MHC coding sequences identified in our samples. Expression was estimated as Fragments Per Kilobase of transcript length per Million reads mapped (FPKM); in the analysis of all seven species we assumed mapping rate of 0.895 as this was the average for the four species with the available genome assemblies.

### Gene tree – species tree reconciliation

Gene tree – species tree reconciliation was performed in Notung v. 3.0_25 beta (Durand et al. 2006) in an attempt to infer the number of duplications and gene losses and identify clusters of orthologs in MHC-I. The analysis was performed on two datasets that used different gene trees: (i) G, and (ii) G+T. Notung analyses used ML trees constructed using protein sequences with RAxML-ng (see above), bootstrap edge weight threshold, which indicates which branches are “weak” set to 50, the cost of duplication and cost of loss were kept at the default values of 1.5 and 1.0. After reconciliation tree was adjusted to further optimise the number of duplications and losses by rearranging “weak” edges.

### Testing for gene conversion

Gene conversion was tested using coding DNA sequences of the G+T dataset after removing sequences assembled by Trinity from short RNAseq reads, as some of them could have been computationally generated chimaeras that may produce a false recombination signal. The analyses were performed separately for MHC-I and MHC-I-like genes, because high sequence divergence between them makes conversion unlikely and because including numerous sequences would reduce the power of the test by inflating the number of pairwise comparisons that would necessitate stringent correction for multiple testing. MHC-I-like sequences from all species were analysed together due to their relatively small number. MHC-I sequences from each species were analysed separately as we were mostly interested in species-specific gene conversion events. The analyses were performed using geneconv v. 1.81a (Padidam et al. 1999) and were run both allowing no mismatches within conversion tracts (*g* = 0) and using the more permissive setting allowing for some mismatches (*g* = 1) and thus enabling detection of older conversion events.

### Selective pressures acting on MHC-I and MHC-I-like genes

Tests of selective pressures were performed applying methods implemented in HyPhy v. 2.5.8 (Kosakovsky Pond et al. 2020), using protein-guided codon alignment of the G+T dataset obtained with DECIPHER:: AlignTranslation() and approximately ML phylogeny inferred with FastTree v. 2.1.11 (Price et al. 2010). RELAX (Wertheim et al. 2015) was used to test whether a change in the strength of selection occurred (i) along the long branches separating MHC-I-like1 and MHC-I-like2 from other sequences in MHC-I phylogeny, (ii) within MHC-I-like1 and MHC-I-like2 clusters. FUBAR (Murrell et al. 2013) was used to identify codons under positive/purifying selection, separately for MHC-I, MHC-I-like1 and MHC-I-like2. FUBAR analysis was limited to α1–α3 domains.

### Structural modelling

Genomic sequences of MHC-I-like1 molecules were obtained for *L. vulgaris* and *P. waltl*, while MHC-I-like2 sequences were identified in *L. helveticus*, *L. vulgaris*, and *T. cristatus*. A putative *P. waltl* MHC-I-like2 locus was also identified on a different chromosome than the core MHC region. This locus contained an ORF spanning 20 exons, several of which appeared to encode domains atypical of MHC molecules. Given its likely chimeric and non-functional nature, this gene was excluded from structural modelling. Signal peptides were predicted using SignalP 6.0 (Teufel et al. 2021) and removed prior to structural modelling. MHC-I-like2 molecules contained short N-terminal sequences resembling leader peptides; however, not confidently identified by SignalP. Therefore, these regions were manually trimmed based on sequence alignment with classical MHC-I molecules. Genomic sequences of all MHC-I-like molecules and one putative classical molecule per species (*L. helveticus*: *Lihe-01*, *L. vulgaris*: *Livu-03*, *T. cristatus*: *Trcr-01*, *P. waltl*: *Plwa-01*), together with corresponding β2m, were modelled with the Alphafold3 Server (Abramson et al. 2024; default settings, access April 2025). Structures were visualized with UCSF ChimeraX 1.9 (Meng et al. 2023). A search for structural homology was performed with DALI server (Holm 2022).

## Data Access

Raw sequence data produced for this study have been deposited in the European Nucleotide Archive under the ENA Research Project accession PRJEB90989. Sequence alignments and do novo annotation of MHC genes are available under the link: https://tinyurl.com/2p3jb3ka and will be deposited as supporting data at the journal web page.

## Acknowledgements

This study was supported by the Polish National Science Centre OPUS Grant No. UMO-2021/43/B/NZ8/00979 to W.B.

