## Supplementary Materials for "Dynamics of complex genomic regions in giant genomes: MHC evolution in newts"

### Suuplementary Tables

Are in a separate Excel workbook

### Supplementary Figure Legends

**Fig. S1. Phylogenies of MHC-I and MHC-I-like domains α1, α2, and α3.** The RAxML-ng maximum likelihood trees constructed from protein sequences under the JTT + G4 amino-acid substitution model. The trees were rooted with *Andrias* sequence (Gen Bank Acc. AGY55973, not shown). Clades with the minimum bootstrap support of 70% (100 bootstrap replicates) are marked with asterisks. The tree contains sequences from the G+T dataset (as described in Materials and Methods) obtained from PacBio IsoSeq, de novo assembly of RNAseq and protein sequences predicted for manually annotated MHC genes in genome assemblies of *L. helveticus*, *L. vulgaris*, *P. waltl* and *T. cristatus*. Sequences predicted in genomes are marked with numbers indicating the number of coding exons. Inset shows time-calibrated phylogeny of the studied species (Stewart and Wiens 2025). MHC sequences are color-coded according to the species. Maximum relative expression – the maximum fraction of total MHC-I expression (FPKM) attributable to a sequence across all RNAseq libraries from postmetamorphic individuals within a species. N conserved AA – the number of residues important for anchoring the termini of antigenic peptides containing amino acids that are conserved in classical MHC-I of most taxa (tippoint on the right). Note, that in α3 tree a single *T. marmoratus* MHC-I-like2 sequence did not cluster together with the remaining MHC-I-like2 sequences, so the labelled MHC-I-like2 clade does not include this sequence.

**Fig. S2. Expression through ontogeny of MHC-I genes annotated in the genomes.** Relative expression, i.e., the fraction of all RNAseq reads in a library mapping to all MHC-I genes and pseudogenes in the assembly that mapped to a particular gene sequence is shown. Boxplots show medians, interquartile and total ranges. Grey stripes indicate putative nonclassical (MHC-Ib) sequences. Species abbreviations: Lh - *L. helveticus*, Lv - *L. vulgaris*, Pw – *P. waltl*, Tc – *T. cristatus*.

**Fig. S3. Expression of MHC-I sequences in adult tissues.** Relative expression, i.e., FPKM for a particular cluster representative divided by the total MHC-I FPKM in a library. RNAseq libraries from different tissues are indicated with different colors and individuals with different shapes. Species abbreviations: Ia – *I. alpestris*, Lb – *L. boscai*, Lh - *L. helveticus*, Lv - *L. vulgaris*, Pw – *P. waltl*, Tc – *T. cristatus*, Tm - *T. marmoratus.*

**Fig. S4. Position of putative classical and nonclassical MHC-I sequences on phylogeny from Fig. 4.** Putative classical (MHC-Ia) and nonclassical (MHC-Ib) sequences were identified as described in the text and their branches are marked red and blue, respectively. The remaining sequences which status is unclear as well as MHC-I-like sequences are marked in gray.

**Fig. S5. Polymorphism of MHC-I genes annotated in the genomes.** Bar heights show polymorphism of each gene expressed relative to the most highly polymorphic gene in the species; MHC-I-like genes are not shown as reliable polymorphism estimates were not available. Polymorphism was estimated based on mapping amplicon variants detected in population samples to genes, as described in the text. Maximum relative expression – the maximum fraction of total MHC-I expression (FPKM) attributable to a sequence across all RNAseq libraries from postmetamorphic individuals within a species. Species abbreviations: Lh - *L. helveticus*, Lv - *L. vulgaris*, Pw – *P. waltl*, Tc – *T. cristatus.*

**Fig. S6. Top views of α1-α2 domains of a) MHC-I-like1 and b) MHC-I-like2 molecules**, generated with Alphafold3. Left hand-side structures: ribbon models of MHC-I-like molecules (magenta – top conformation of MHC-I-like1, blue – alternative MHC-I-like1 conformation, brown - top conformation of MHC-I-like2), overlaid on classical molecules of respective species. Middle structure: Coulombic electrostatic potential mapped on the molecular surface, with red for negative and blue for positive potential. Right hand-side structures: lipophilicity potential mapped on the molecular surface, with cyan most hydrophilic and goldenrod most lipophilic.

**Fig. S7. The relationship between MHC-I and TAP (TAP1 + TAP2) diversity in *Triturus* newts.** Estimates of phylogenetic alpha (within-individual) and gamma (within-population ) diversity for 30 salamander species are from Palomar et al. (2021). There is a tendency for *Triturus* species (color-coded) to show low TAP diversity and little evidence for a correlation between MHC-I and TAP diversity, as opposed to the remaining 23 species representing 15 genera and six families.

**Fig. S1.**

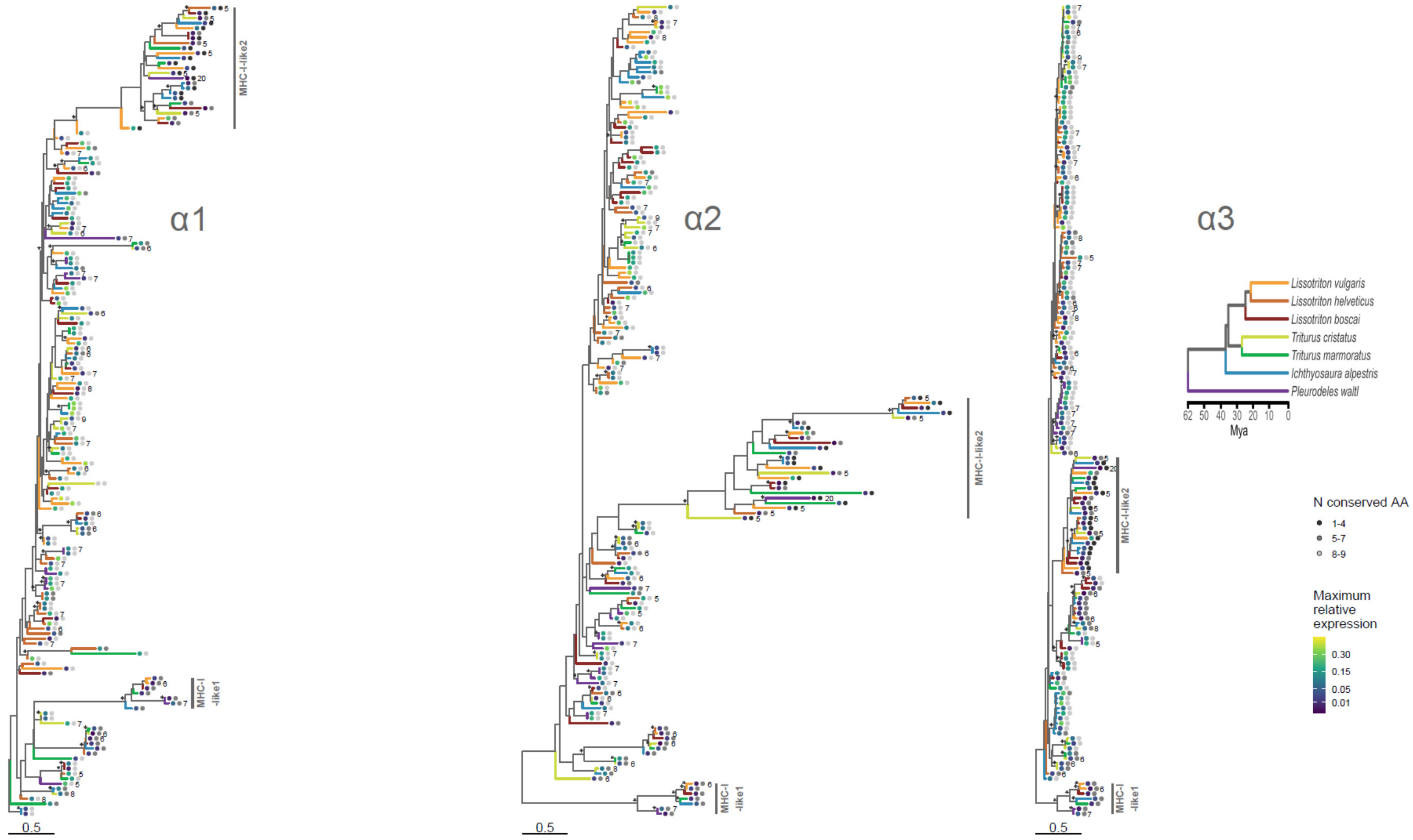
α1
α2
α3
MHC-I-like2
MHC-I -like1
0.5
Lissotriton vulgaris
Lissotriton helveticus
Lissotriton boscai
Triturus cristatus
Triturus marmoratus
Ichthyosaura alpestris
Pleurodeles waltl
62 50 40 30 20 10 0
Mya
N conserved AA
1-4
5-7
8-9
Maximum relative expression
0.30
0.15
0.05
0.01

**Fig. S2.**

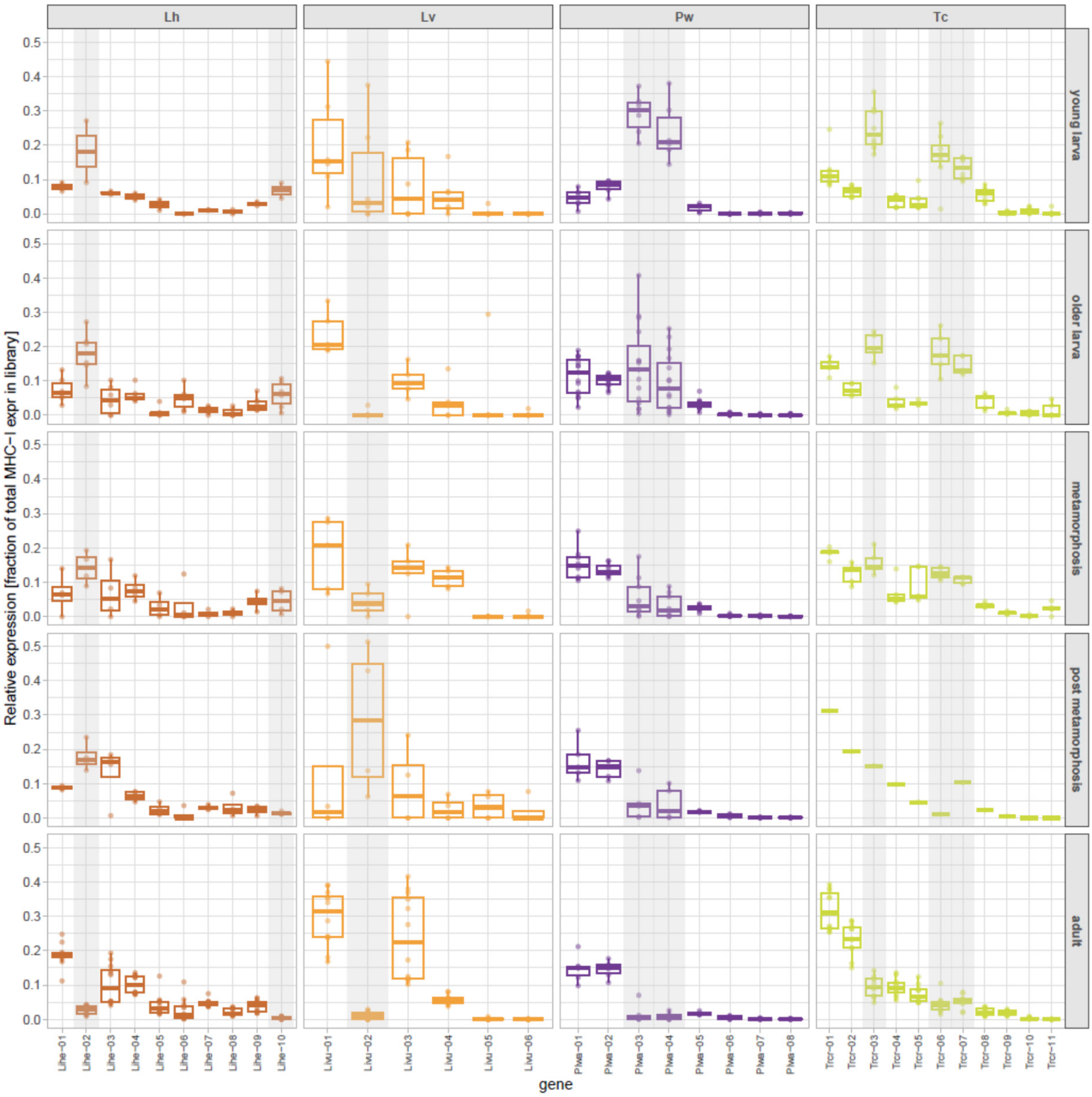
Lh
Lv
Pw
Tc
young larva
older larva
metamorphosis
post metamorphosis
adult
Relative expression [fraction of total MHC−I expr in library]
0.0
0.1
0.2
0.3
0.4
0.5
gene
Lihe−01
Lihe−02
Lihe−03
Lihe−04
Lihe−05
Lihe−06
Lihe−07
Lihe−08
Lihe−09
Lihe−10
Livu−01
Livu−02
Livu−03
Livu−04
Livu−05
Livu−06
Plwa−01
Plwa−02
Plwa−03
Plwa−04
Plwa−05
Plwa−06
Plwa−07
Plwa−08
Trcr−01
Trcr−02
Trcr−03
Trcr−04
Trcr−05
Trcr−06
Trcr−07
Trcr−08
Trcr−09
Trcr−10
Trcr−11

**Fig. S3.**

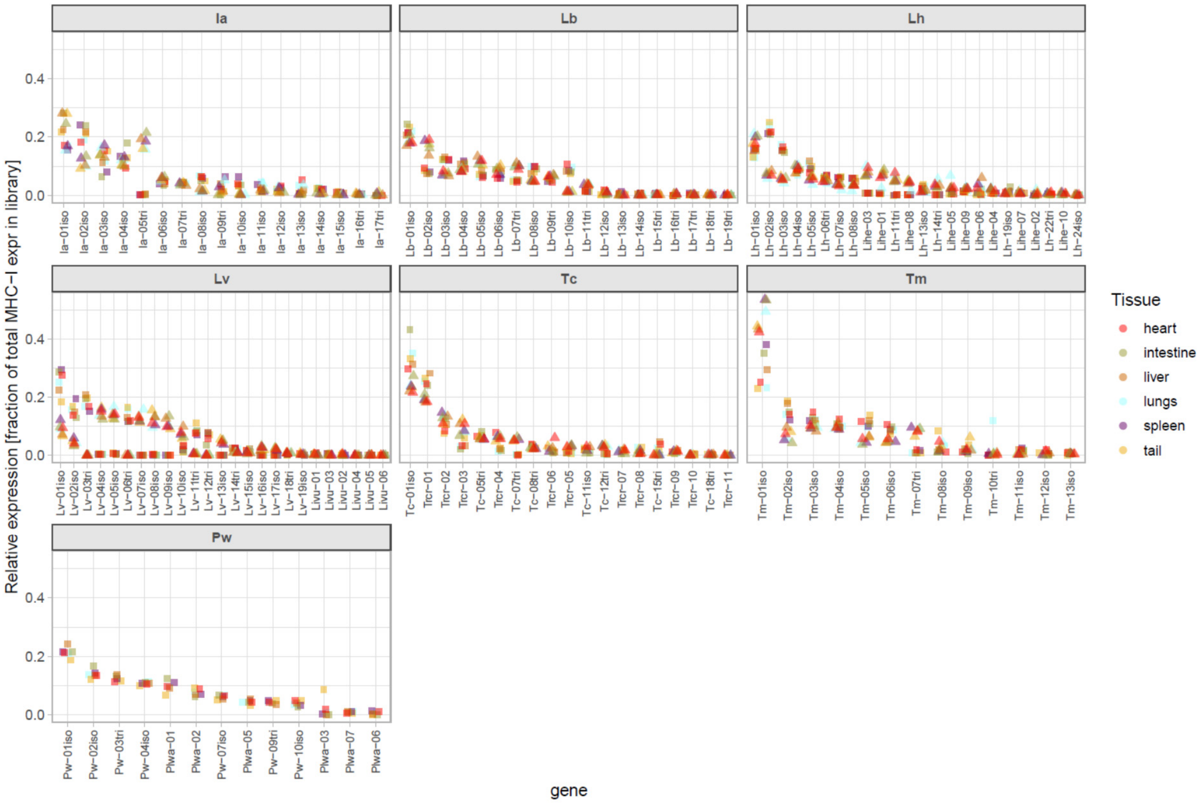
Ia
Lb
Lh
Lv
Tc
Tm
Pw
Relative expression [fraction of total MHC−I expr in library]
0.0
0.2
0.4
gene
Ia−01iso Ia−02iso Ia−03iso Ia−04iso Ia−05tri Ia−06iso Ia−07tri Ia−08iso Ia−09tri Ia−10iso Ia−11iso Ia−12iso Ia−13iso Ia−14iso Ia−15iso Ia−16tri Ia−17tri
Lb−01iso Lb−02iso Lb−03iso Lb−04iso Lb−05iso Lb−06iso Lb−07tri Lb−08iso Lb−09tri Lb−10iso Lb−11tri Lb−12iso Lb−13iso Lb−14iso Lb−15tri Lb−16tri Lb−17tri Lb−18tri Lb−19tri
Lh−01iso Lh−02iso Lh−03iso Lh−04iso Lh−05iso Lh−06tri Lh−07iso Lh−08iso Lihe−03 Lihe−01 Lh−11tri Lihe−08 Lh−13iso Lh−14tri Lihe−05 Lihe−09 Lihe−06 Lihe−04 Lh−19iso Lihe−07 Lihe−02 Lh−22tri Lihe−10 Lh−24iso
Lv−01iso Lv−02iso Lv−03tri Lv−04iso Lv−05iso Lv−06tri Lv−07iso Lv−08iso Lv−09iso Lv−10iso Lv−11tri Lv−12tri Lv−13iso Lv−14tri Lv−15iso Lv−16iso Lv−17iso Lv−18tri Lv−19iso Livu−01 Livu−03 Livu−02 Livu−04 Livu−05 Livu−06
Tc−01iso Trcr−01 Trcr−02 Trcr−03 Tc−05tri Trcr−04 Tc−07tri Tc−08tri Trcr−06 Trcr−05 Tc−11iso Tc−12tri Trcr−07 Trcr−08 Tc−15tri Trcr−09 Trcr−10 Tc−18tri Trcr−11
Tm−01iso Tm−02iso Tm−03iso Tm−04iso Tm−05iso Tm−06iso Tm−07tri Tm−08iso Tm−09iso Tm−10tri Tm−11iso Tm−12iso Tm−13iso
Pw−01iso Pw−02iso Pw−03tri Pw−04iso Plwa−01 Plwa−02 Pw−07iso Plwa−05 Pw−09tri Pw−10iso Plwa−03 Plwa−07 Plwa−06
Tissue
heart
intestine
liver
lungs
spleen
tail

**Fig. S4.**

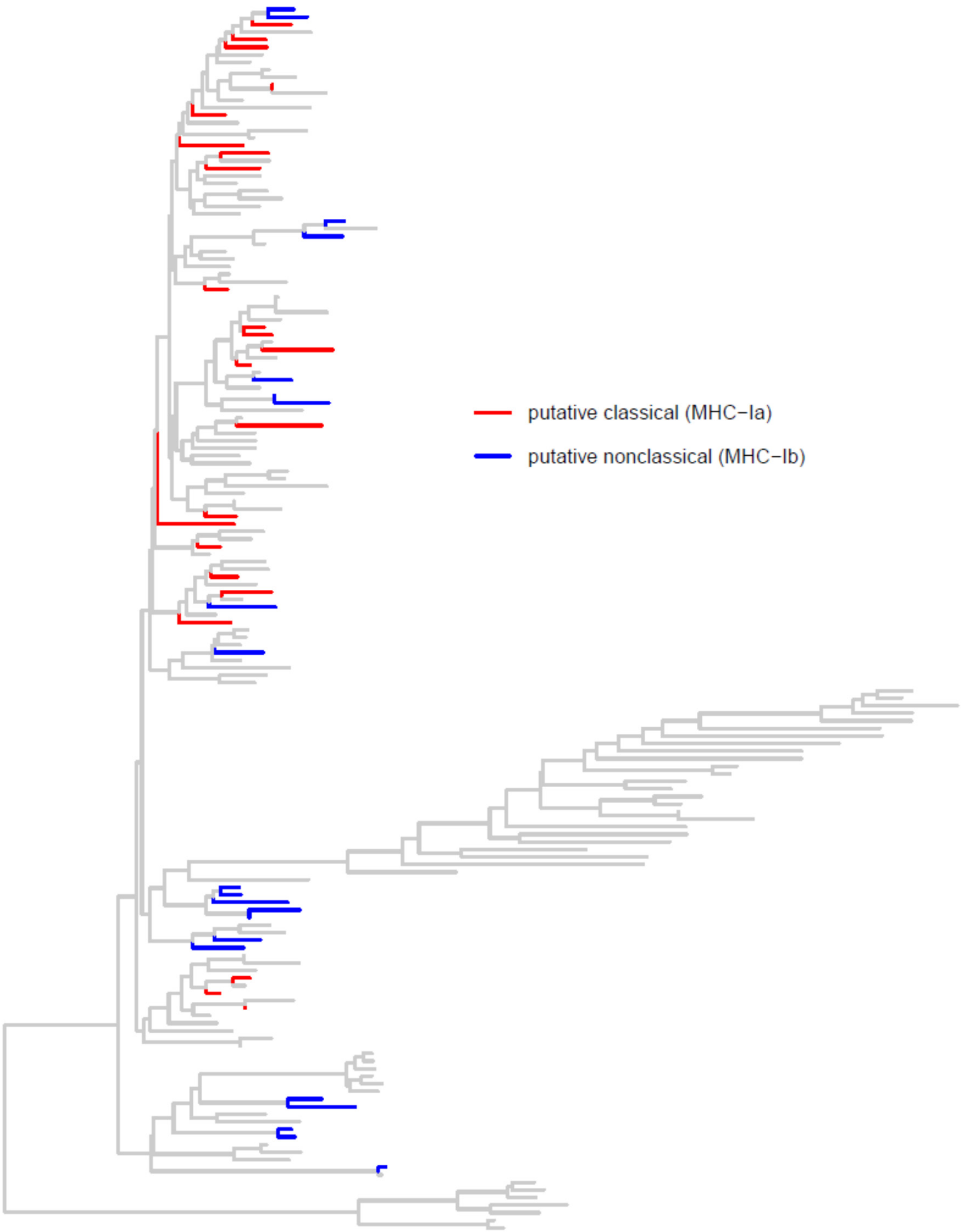

**Fig. S5.**

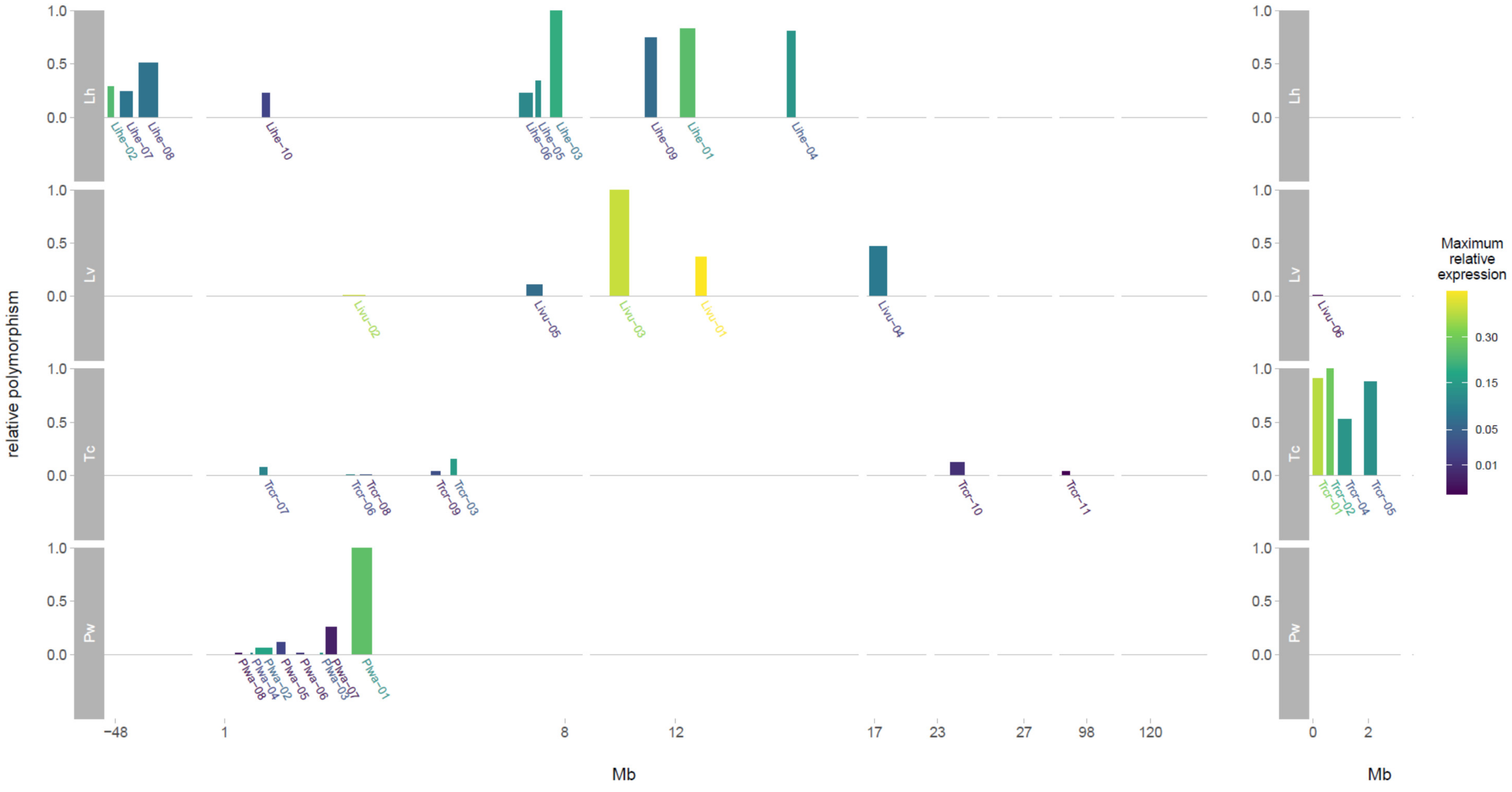
relative polymorphism
Lh
Lv
Tc
Pw
1.0
0.5
0.0
Lihe−02
Lihe−07
Lihe−08
Lihe−10
Lihe−06
Lihe−05
Lihe−03
Lihe−09
Lihe−01
Lihe−04
Livu−02
Livu−05
Livu−03
Livu−01
Livu−04
Livu−06
Trcr−07
Trcr−06
Trcr−08
Trcr−09
Trcr−03
Trcr−10
Trcr−11
Trcr−01
Trcr−02
Trcr−04
Trcr−05
Plwa−08
Plwa−04
Plwa−02
Plwa−05
Plwa−06
Plwa−03
Plwa−07
Plwa−01
−48
1
8
12
17
23
27
98
120
0
2
Mb
Mb
Maximum relative expression
0.30
0.15
0.05
0.01

**Fig. S6.**

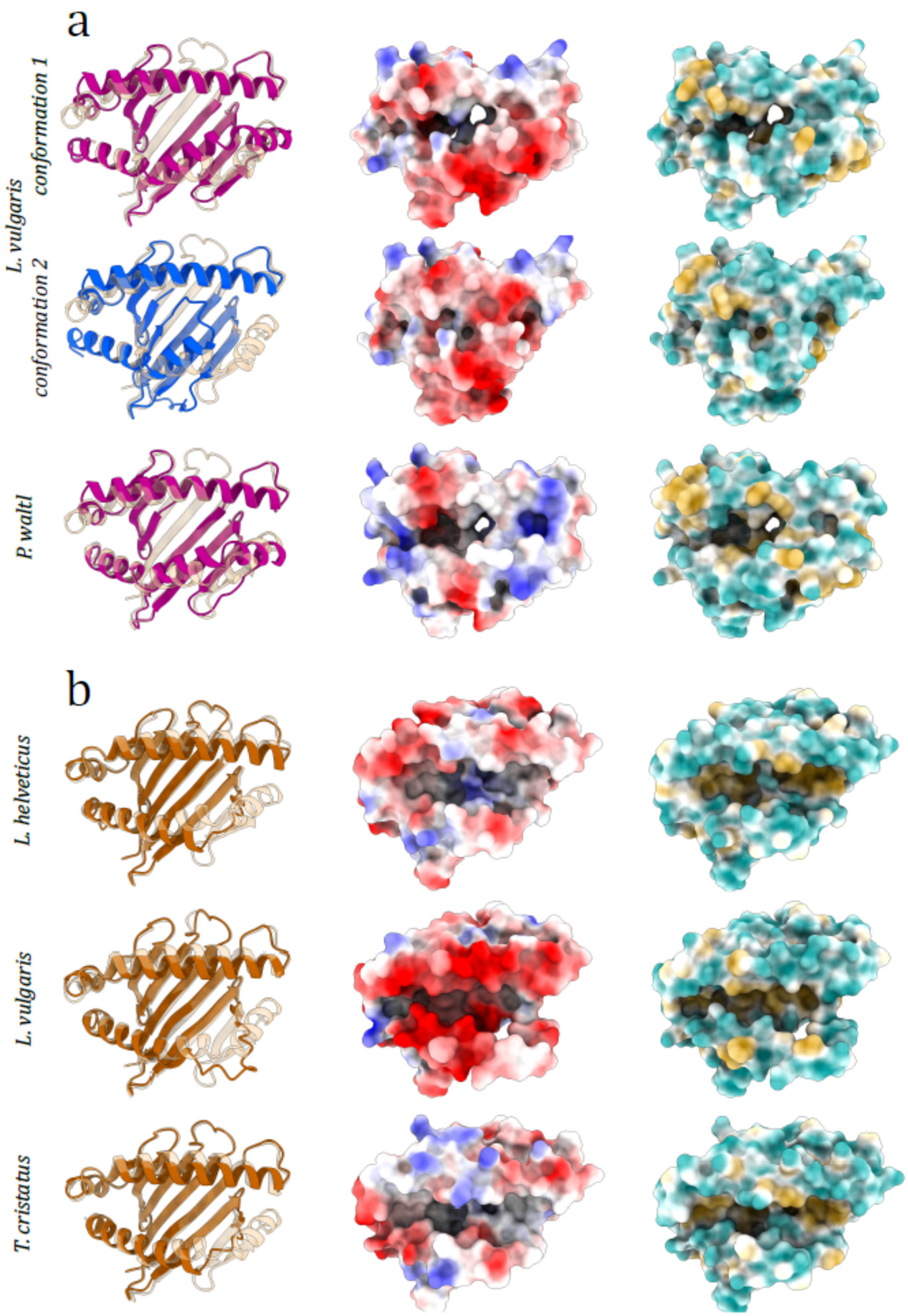
a
L. vulgaris
conformation 1
conformation 2
P. waltl
b
L. helveticus
L. vulgaris
T. cristatus

**Fig. S7.**

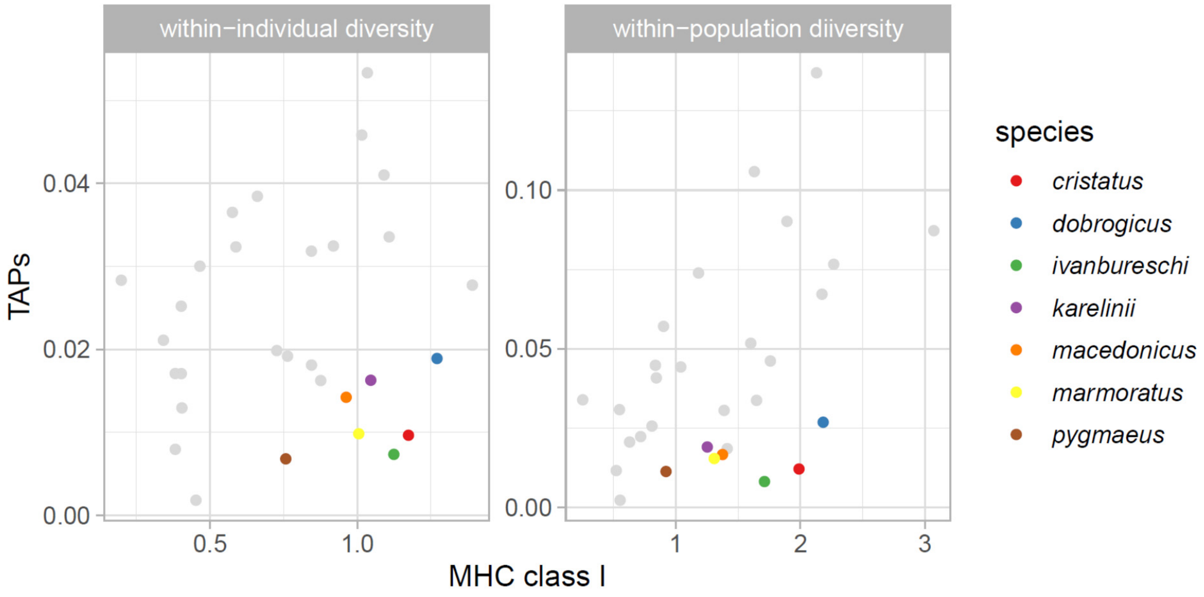
within−individual diversity
within−population diiversity
TAPs
0.04
0.02
0.00
0.10
0.05
0.00
0.5
1.0
1
2
3
MHC class I
species
cristatus
dobrogicus
ivanbureschi
karelinii
macedonicus
marmoratus
pygmaeus
